# Immunotherapy efficacy in colorectal cancer is dependent on activation of a microbial-metabolite-immune circuit

**DOI:** 10.1101/2020.02.26.965533

**Authors:** Lukas F. Mager, Regula Burkhard, Noah C.A. Cooke, Kirsty Brown, Hena Ramay, Seungil Paik, John Stagg, Ryan A. Groves, Marco Gallo, Ian A. Lewis, Markus B. Geuking, Kathy D. McCoy

**Author notes:** Corresponding authors: Kathy D. McCoy and Lukas F. Mager.

## Abstract

Cancer is a leading cause of death globally. Checkpoint blockade therapies offer a promising treatment for many cancers, but have been ineffective for colorectal cancers. Previous studies have shown a dependency of immunotherapies on the microbiota. Consequently, we hypothesized that specific gut bacteria promote immunotherapy for colorectal cancer. We identify three commensal bacteria and a microbial metabolite, inosine, that enhance the efficacy of immune checkpoint blockade therapy in colorectal cancer. We show that inosine interacts with the adenosine A_2A_ receptor on T cells resulting in intestinal Th1 cell differentiation. Decreased gut barrier function induced by immunotherapy increased the translocation of bacterial metabolites and promoted cancer protective Th1 cell activation. This microbial-metabolite-immune circuit provides a mechanism for a new class of bacteria-enhanced checkpoint blockade therapies. The efficacy of this mechanism differs among colorectal cancer subtypes and highlights the strengths as well as potential limitations of this novel bacterial co-therapy for cancer.

## Introduction

Colorectal cancer (CRC) is the second and third most common malignancy in Western countries in women and men, respectively (Ferlay et al., 2015). In addition to genetic aberrations, which are essential for the development of CRC, other disease-contributing factors have been identified. These include the microbiota and inflammation, whereby inflammation can drive or inhibit CRC development. Interferon (IFN)-γ producing T helper type 1 (Th1) cells are known to be protective (Mager et al., 2016; Mlecnik et al., 2016; Wang et al., 2015), whereas interleukin (IL)-17-producing Th17 cells promote CRC development (Galon et al., 2006; Grivennikov et al., 2012; Le Gouvello et al., 2008). In fact, the impact of the immune system is so potent that immune cell infiltration in the tumor is a superior prognostic factor compared to the classical tumor-lymph nodes-metastasis (TNM) system in CRC (Anitei et al., 2014; Mlecnik et al., 2016). Similarly, the microbiota also impacts on CRC progression(Arthur et al., 2012; Dejea et al., 2018) and may even alter the efficacy of chemotherapeutics (Iida et al., 2013; Viaud et al., 2013).

Immune checkpoint blockade (ICB) therapy is an efficient anti-cancer strategy that utilizes the therapeutic potential of the immune system. Most notably, ICB inhibitors targeting cytotoxic T-lymphocyte-associated antigen 4 (CTLA-4), programmed cell death protein 1 (PD-1), or its ligand (PD-L1) have shown great success in the treatment of various cancers, including melanoma, renal cell carcinoma, and non-small cell lung cancer (Brahmer et al., 2012; Hodi et al., 2010). More recently, seminal work has shown that the efficacy of ICB therapy is dependent on the presence of certain ICB-promoting gut bacteria (Routy et al., 2018; Sivan et al., 2015; Vetizou et al., 2015).

Despite these exciting advances, ICB therapy efficacy in CRC has been disappointing (Brahmer et al., 2012), with only 5-10% of all CRC patients responding (Le et al., 2017). Moreover, the detailed molecular mechanisms through which bacteria enhance the efficacy of ICB therapies remains unclear. Here, we identified three bacterial species that promote ICB efficacy in CRC and identified inosine as a critical bacterial metabolite that promoted differentiation of Th1-mediated anti-tumor immunity.

## Results

### ICB therapy efficacy depends on the gut microbiota

We first questioned whether ICB therapy efficacy in CRC is dependent on the microbiota. Heterotopic MC38 colorectal cancers were implanted into germ-free (GF) and specific pathogen free (SPF) mice and, upon palpable tumor development, ICB therapy was initiated, which led to smaller tumors in SPF animals (Figure S1A and B). Moreover, intratumoral and splenic CD4^+^ and CD8^+^ T cell activation and proliferation were markedly increased in SPF animals (Figure S1C - N). To ensure this was not merely a reflection of the immature immune system of GF mice, we also assessed the effect and ICB therapy in antibiotic-treated SPF mice (Figure S1O). Compared to control treatment, broad spectrum antibiotics also reduced ICB therapy efficacy in tumor-bearing SPF mice (Figure S1P - T). These results indicated that ICB efficacy is enhanced in the presence of microbes, corroborating previous reports with other tumor types (Vetizou et al., 2015).

### Identification of ICB-promoting bacteria in CRC

Clinically, ICB therapies are notoriously ineffective in most CRC cases (Le et al., 2015) and heterotopic tumors may not adequately model the spatially close interactions between the gut microbiota and local immunity in intestinal tumors. We therefore employed a more physiological model of CRC to investigate the interactions between the microbiota and immunity in the context of ICB therapy. Intestinal tumors were induced using azoxymethane (AOM) and dextran sulfate sodium (DSS) in SPF animals. Following tumor development, we evaluated the ability of ICB therapy to induce anti-tumor immunity (Figure 1A). Notably, ICB therapy led to smaller and fewer tumors (Figure 1B and C), reduced cancer stem cell numbers (Figure 1D), increased immune cell infiltration into the tumors (Figure 1E), and increased CD8^+^ T cell frequencies in the tumor draining lymph node together with increased splenic CD4^+^ and CD8^+^ T cell activation (Figure 1F-H). In this model, anti-CTLA-4 tumoricidal effects were greater than those induced by anti-PD-L1 treatment when using the same antibody dose. In order to identify potentially beneficial tumor-associated bacteria, we performed 16S rRNA gene V4 region amplicon sequencing of genomic DNA isolated from homogenized tumors as well as anaerobic culture of homogenized tumor tissue. Microbial sequencing revealed that the tumor-associated bacterial community composition of ICB-treated tumors differed from that of control-treated tumors (Figure S2A and Figure 1I). Furthermore, we were able to culture twenty-one different bacterial species from tumor tissues. Notably, seven of these cultured bacteria were found only in the ICB-treated group, whereas four were found only in the control group (Figure 1J).

**Figure 1:**
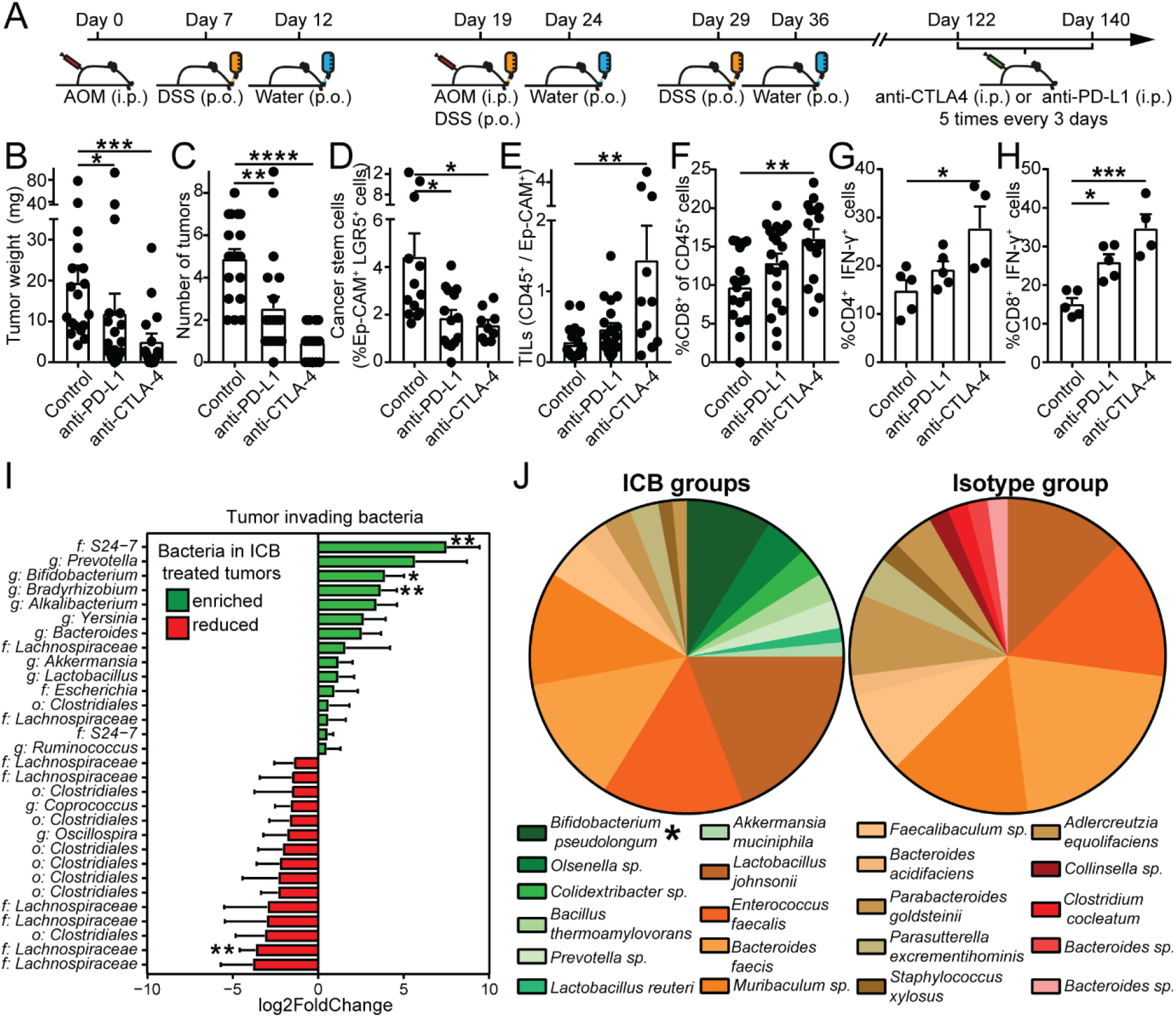
Immune cell and microbial dynamics upon ICB therapy in AOM/DSS tumors. (A) Overview of the experimental setup of AOM/DSS-induced CRC and ICB treatment. (B) Tumor weight, (C) number of tumors, (D) Ep-CAM^+^, LGR5^+^ cancer stem cells and (E) tumor-infiltrating leukocytes (TILs) 140 days post induction in animals treated with isotype-, anti-PD-L1 or anti-CTLA-4 antibodies. (F) CD8^+^ T cell frequencies in the tumor draining lymph node at day 140. Splenic IFN-γ^+^ production in (G) CD4^+^ or (H) CD8^+^ T cells. (I) 16S rRNA gene V4 region amplicon sequencing to identify bacteria in tumor tissue. Bacteria enriched or reduced in tumors of anti-PD-L1/anti-CTLA-4 compared to isotype treated animals are shown in green or red, respectively. (J) Bacteria cultured from homogenized tumors under anaerobe conditions from anti-PD-L1/anti-CTLA-4 (ICB groups) or isotype (Isotype group) treated animals. Bacteria depicted as green or red could only be cultured in ICB groups or Isotype group, respectively. Bacteria depicted as brown were present in both groups. Data are (B-H) mean ± SEM or (I) mean ±lfcSE (logfoldchangeStandard Error) and pooled from three individual experiments. (B-F) *n* =16-20 mice/group, (g and h) *n* = 4-5 mice/group. *, *P* < 0.05; **, *P* < 0.01; ***, *P* < 0.001; ****, *P* < 0.0001. Sea also Figures: S1 and S2

Interestingly, *Akkermansia muciniphilia,* which was recently identified to enhance the efficacy of anti-PD-L1 and anti-PD-1 treatments in lung and kidney cancers (Routy et al., 2018), was one of the seven bacteria cultured only from ICB-treated tumors. We also performed 16S rRNA gene V4 amplicon sequencing of fecal samples from control and ICB groups but found no significant differences in microbiota composition (Figure S2B, S2C and Table S1), indicating that the tumor-associated bacterial communities provided a better source for identification of ICB-promoting bacteria in CRC.

To address whether the bacteria that were found to be enriched in the ICB-treated tumors were able to boost the efficacy of ICB therapy, we selected five of the isolated culturable bacterial species for monocolonization of GF mice. Monocolonized or GF mice were injected with MC38 tumor cells, treated with anti-CTLA-4 upon palpable tumor development and assessed for effects on tumor growth and anti-tumor immunity (Figure 2A). We chose the heterotopic model of CRC for this approach as the development of orthotopic CRC is severely reduced in animals with a limited microbiota (Schwabe and Jobin, 2013). Of the five bacteria tested, monocolonization with *Bifidobacterium pseudolongum* (*B.p.*), *Lactobacillus johnsonii* (*L.j.*), and *Olsenella sp*. (*O.sp*.) significantly enhanced the efficacy of anti-CTLA-4 treatment compared to GF mice or mice monocolonized with *Colidextrbacter sp.* (*C.sp*.) or *Prevotella sp*. (*P.sp*.) (Figure 2B-E). In addition, CD4^+^ and CD8^+^ T cell activation and proliferation were substantially increased in the tumors of *B.p.*, *L.j.*, and *O.sp.* monocolonized animals (Figure 2F-I). The isolated ICB-promoting *B.p. strain* also improved the efficacy of anti-PD-L1 treatment in the MC38 heterotopic tumor model compared to the *C.sp.* control strain (Figure S3), albeit to a lower extent than observed for anti-CTLA-4 treatment (at the same dose), which is similar to our observations in the AOM/DSS model. Due to its greater observed efficacy, we performed all subsequent mechanistic studies using anti-CTLA-4 treatment. Of note, anti-tumor immunity was dependent on anti-CTLA-4 or anti-PD-L1 co-therapy as monocolonization with *B.p.* alone was not able to reduce tumor growth (Figure S4A-D) or induce anti-tumor immunity (Figure S4E-J), similar to previous studies with other ICB-promoting bacteria(Routy et al., 2018; Vetizou et al., 2015).

**Figure 2:**
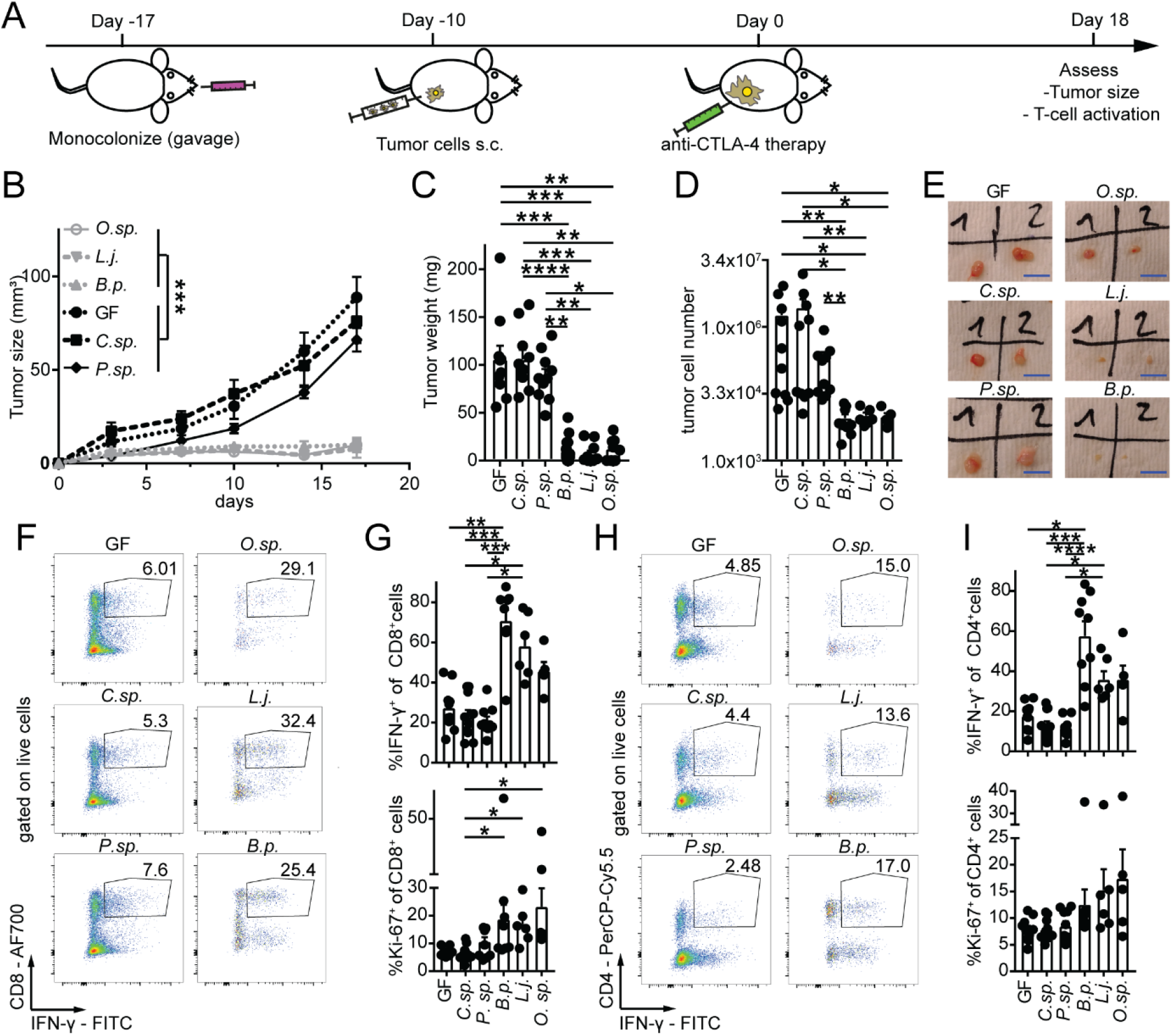
Individual bacterial species boost ICB therapy. (A) Schematic of the experimental setup to assess the effect of individual bacteria on anti-CTLA-4 therapy efficacy. (B) Tumor growth, (C) tumor weight, (D) live tumor cells and (E) representative pictures of tumors are shown at day 18. Scale bars: 1cm. (F) representative plots and (G) quantification of IFN-γ^+^ and Ki-67^+^, CD8^+^ T cells at day 18 in the tumor tissue. (H and I) same as (F and G), but for CD4^+^ T cells. Data are mean ± SEM and pooled from three individual experiments (*n* = 8-15 mice/group). *, *P* < 0.05; **, *P* < 0.01; ***, *P* < 0.001; ****, *P* < 0.0001. See also Figures: S3, S4 and S6

### Induction of Th1 immunity through ICB promoting bacteria

To investigate the mechanism by which the identified bacteria enhanced ICB therapy we selected *B.p.* as a representative of the beneficial bacteria since it appeared to have the strongest ICB-promoting effect. GF or *C.sp*. monocolonized served as negative controls. Previous studies revealed the ability of some bacteria to accumulate in the tumor environment where they locally stimulate the immune system and kill tumor cells through toxic metabolites (Zheng et al., 2018). Although bacteria were abundantly present in the feces of *B.p.* and *C.sp*. monocolonized mice, we were unable to detect bacteria or amplify 16S rDNA from heterotopic tumors of these mice (Figure S5), indicating that the beneficial effect of bacteria in this model does not require bacteria to reside within the tumor itself. Compared to GF or *C.sp*. monocolonized mice, *B.p.* monocolonization induced a significant increase in expression of the Th1 master transcription regulator T-bet in small intestinal lamina propria CD4^+^ T cells. Similarly, albeit to a lower extent, *B.p.* induced T-bet expression in CD4^+^ T cells in the mesenteric lymph nodes (MLN), but not in the spleen (Figure 3A-G). Intriguingly, *B.p.* did not activate the effector function of Th1 cells as T-bet^+-^IFN-γ ^+^ double positive cells did not differ between *B.p*, *C.sp*. or GF groups in any of the tissues assessed. Taken together, in the absence of tumors and ICB therapy, *B.p.* alone promoted Th1 transcriptional differentiation without increasing effector function locally in the gut and draining lymph nodes but not systemically. While *B.p.* had no effect on other CD4^+^ T cell subsets in the small intestine, it also increased CD8^+^T-bet^+^ T cells (Figure S6A-S6E). Moreover, *B.p.* had minimal impact on Th17 and Treg cells in the MLN and spleen (Figure S6F-S6O).

**Figure 3:**
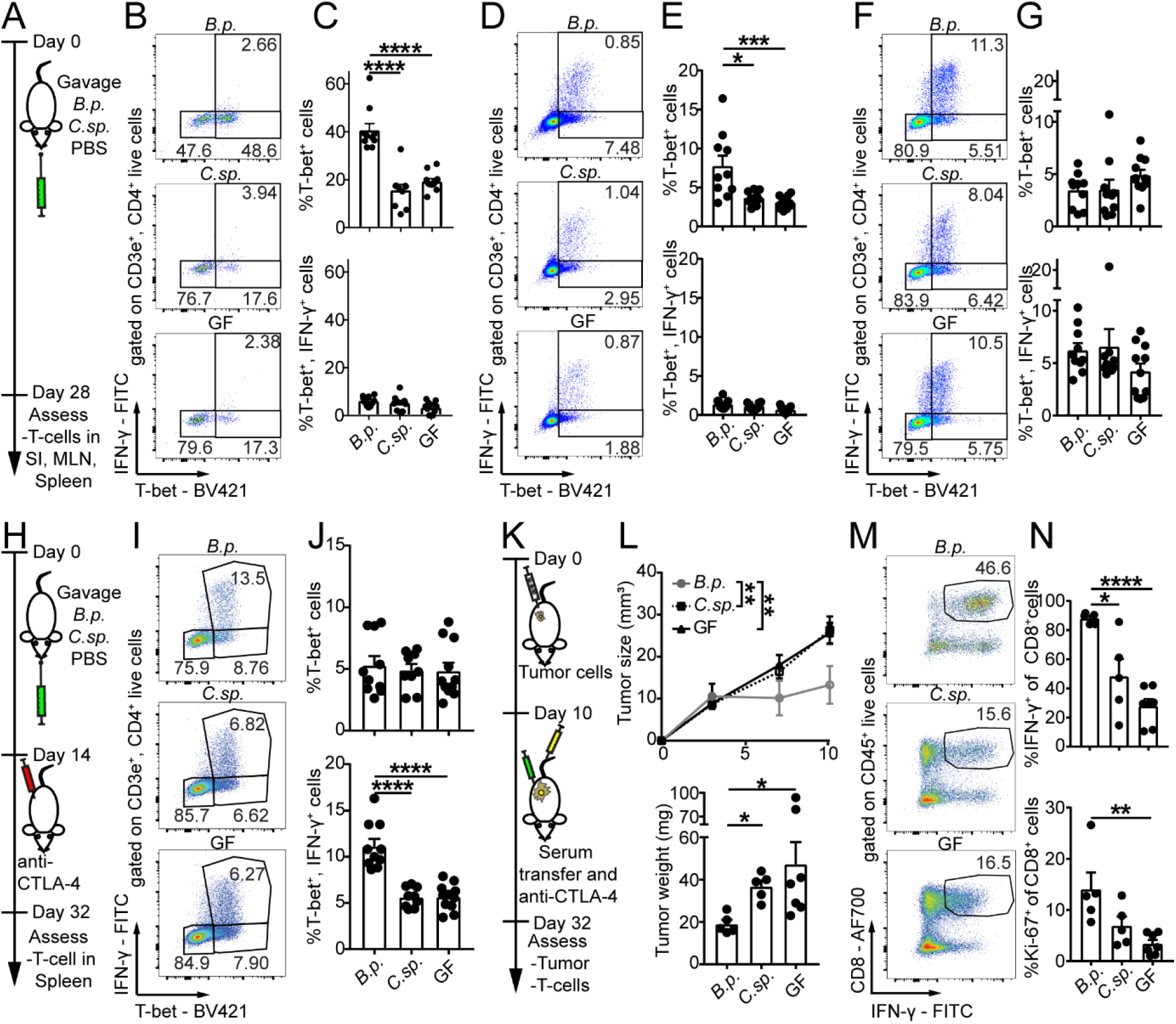
Effect of *B.p.,* anti-CTLA-4 and *B.p.* conditioned serum on T cell differentiation and activation. (A, H and K) Schematic of the experimental setups. (B) representative plots and (C) quantification of T-bet^+^ and T-bet^+^, IFN-γ^+^ events of CD3^+^, CD4^+^ cells in the small intestine (SI) in the presence of indicated bacteria at day 28. (D and E) same as (B and C), but in the mesenteric lymph node (MLN). (F and G) same as (B and C,) but in the spleen. (I) representative plots and (J) quantification of T-bet^+^ and T-bet^+^, IFN-γ^+^ events of CD3^+^, CD4^+^ T cells in the spleen in the presence of indicated bacteria and anti-CTLA-4 treatment at day 32. (L) Tumor growth and weight are shown 32 days after MC38 tumor challenge and subsequent serum transfer as well as anti-CTLA-4 treatment. (m) representative plots and (n) quantification of intratumoral IFN-γ ^+^ or Ki-67^+^CD8^+^ T cells. Data are mean ± SEM and pooled from two individual experiments (A-J) *n* = 10-11 mice/group, (K-N). *n* = 5-7 mice/group. *, *P* < 0.05; **, *P* < 0.01; ***, *P* < 0.001; ****, *P* < 0.0001. See also Figures: S5, S6, S7 and S8

### Systemic effect of ICB promoting bacteria

Since *B.p.* alone promoted only local and not systemic Th1 differentiation during homeostasis, we next asked whether the combination of *B.p.* monocolonization and anti-CTLA-4 therapy (in the absence of a tumor) would induce systemic Th1 activation. Indeed, when combined with anti-CTLA-4, *B.p.* was able to significantly enhance splenic Th1 cell activation and effector function as evidenced by IFN-γ production compared to *C.sp*. monocolonized or GF animals (Figure 3H-J, Figure S6P and S6Q). We concluded that *B.p.* induces Th1 differentiation and, together with anti-CTLA-4, activation of Th1 T cells. Interestingly, a recently defined consortium of eleven bacteria was found to induce IFN-γ production preferentially in CD8^+^ T cells and promote anti-tumor immunity in the absence of immunotherapy (Tanoue et al., 2019). In contrast, *B.p*.-induced IFN-γ production in both CD4^+^ and CD8^+^ T cells (Figure S6R), and ICB treatment was required for tumoricidal function. We were intrigued by the ability of *B.p.* to induce Th1 transcriptional differentiation during homeostasis versus activation of effector function following ICB treatment. Gastrointestinal inflammation is a common immune-related adverse effect of anti-CTLA-4 treatment (Hodi et al., 2010) and we reasoned that this may be due to alterations in gut barrier integrity. Indeed, animals treated with anti-CTLA-4 had increased systemic serum anti-commensal antibodies, particularly Th1-associated IgG2b (Germann et al., 1995), and reduced small intestinal transepithelial electrical resistance compared to controls (Figure S7A and S7B). Although anti-CTLA-4 treatment caused impairment of intestinal barrier integrity it did not induce local or systemic inflammation (Figure S7C and S7D). Since bacteria did not accumulate in the (heterotopic) tumors and anti-CTLA-4 reduced the integrity of the gut barrier, we hypothesized that increased systemic translocation of metabolites may be responsible for the systemic effect of *B.p.* during ICB therapy. To address this, we collected serum from tumor-bearing GF, *B.p.* or *C.sp*. monocolonized mice treated with anti-CTLA-4 (see Figure 2A) and transferred it concomitantly with anti-CTLA-4 into GF MC38 tumor-bearing mice. Remarkably, serum from CTLA-4-treated *B.p.* monocolonized mice, but not from GF or *C.sp*. monocolonized mice, was sufficient to reduce tumor growth and elicit strong anti-tumor immunity in the tumor and spleen of GF mice (Figure 3K-N, Figure S8A-S8F). In sum, these data show that soluble factors derived from or induced by *B.p.* were responsible for the observed ICB-promoting effects.

### Molecular mechanism underlying Th1 immune cell differentiation

In order to identify putative metabolites that might be responsible for the anti-tumor effects of the transferred serum, we determined the metabolomic profile of the transferred serum samples and identified metabolites that were increased in the serum of mice monocolonized with *B.p.* compared to *C.sp*. or GF mice. Untargeted metabolomics analysis revealed increased levels of several metabolites in sera from *B.p.* compared to *C.sp*. monocolonized and GF mice (Figure 4A, Figure S9A and S9B). Notably, the purine metabolite inosine was the only metabolite that was significantly more abundant (8 to 9-fold) in sera from *B.p.* monocolonized mice compared to sera from *C.sp*. monocolonized or GF mice (Figure 4B). Of note, xanthine and hypoxanthine, degradation products of inosine, were also elevated in the sera of *B.p.* monocolonized mice (Table S2). Analysis of bacterial culture supernatant revealed that *B.p.* produced approximately ten-fold higher amounts of inosine than *C.sp.* cultured under the same culture conditions, revealing that inosine is a bacterial metabolite produced by *B.p.* Of note, *L.j.* produced higher amounts of hypoxanthine, a potent ligand binding to the same receptor as inosine. (Figure S9C). The identity of inosine was confirmed by fragmentation analysis (Figure S9D). To determine physiological inosine levels *in vivo* we next measured inosine concentrations in duodenal, jejunal and cecal contents of *B.p.* monocolonized mice. Inosine concentrations were highest in the duodenum and gradually decreased along the gastrointestinal tract (duodenum 66.13 ±14.23 μM >jejunum 29.26 ± 9.38μM >cecum 0.5 ± 0.05μM; Figure S9E). We also quantified inosine concentrations in the serum of *B.p.* (26.16 ± 3.32μM) and *C.sp.* (3.26 ±1.01μM) monocolonized mice (Figure S9E), verifying our previous results. Furthermore, inosine levels in the serum of SPF mice (4.08 ± 1.12μM) increased significantly following anti-CTLA-4 treatment (11.65 ± 2.09μM) and this was greatly diminished in antibiotic-treated SPF mice (2.03 ± 0.86μM) (Figure S9F). These data indicated that bacterial production in the upper gastrointestinal tract is likely to be the predominant source of systemic inosine in *B.p.* monocolonized mice.

**Figure 4:**
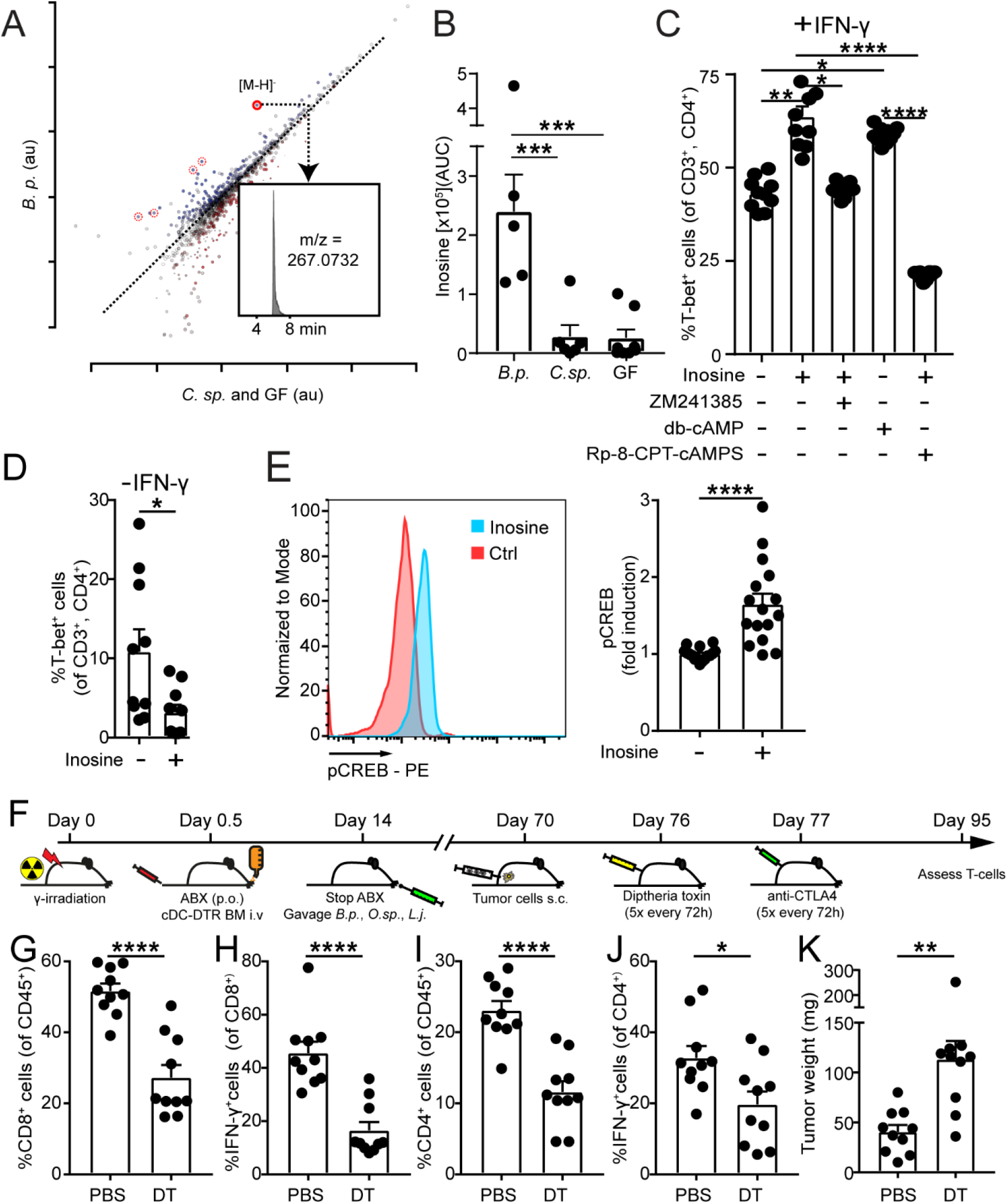
Effect of inosine on T cell differentiation and dependency on classical dendritic cells of ICB therapy efficacy. (A) Scatter plot of untargeted metabolomics data in the serum of anti-CTLA-4 treated, tumor-bearing *B.p.* monocolonized compared to *C.sp*. monocolonized and GF mice. Red circles or dotted red circles depict inosine or inosine fragments/adducts, respectively. Inset shows an extracted ion chromatogram of inosine (B) Intensity of inosine (AUC: area under the curve) in sera shown in panel (A) of this figure. (C) Naïve CD4^+^ T cells were co-cultured with bone marrow derived dendritic cells and IFN-γ. Quantification of T-bet^+^, CD3^+^, CD4^+^ T cells 48 hours after co-culture in the presence or absence of inosine, A_2A_ receptor inhibitor (ZM241385), cell permeable cAMP (db-cAMP) and protein kinase A inhibitor (RP-8-CPT-cAMPS). (D) Same as (C) without IFN-γ. (E) Representative plot and quantification (left and right panel) of phospho-CREB (Ser133) levels in naïve CD4^+^ T cells cultured with anti-CD3/anti-CD28 coated beads for 1 hour in the presence or absence of inosine. (F) Schematic overview of experimental setup to deplete classical dendritic cells during MC38 tumor challenge and anti-CTLA-4 treatment. Intratumoral (G) frequency of CD8^+^ T cells (H) IFN-γ^+^CD8^+^ T cells. (I and J) same as (G and H), but CD4^+^ T cells and (K) tumor weight at day 95. Data are mean ± SEM and pooled from two individual experiments (A-B) *n* = 5-8 samples/group (C-E) 10-15 biological replicates/group (F-K) 10 mice/group. *, *P* < 0.05; **, *P* < 0.01; ***, *P* < 0.001; ****, *P* < 0.0001. See also Figures: S9, S10, S11 and S12

We next investigated whether inosine could directly enhance anti-tumor Th1 cell differentiation. To test this, we co-cultured activated OVA_323-339_ peptide-pulsed bone marrow derived dendritic cells (BMDCs) with naïve OVA_323-339_-specific OT-II CD4^+^ T cells in the presence or absence of inosine. Intriguingly, inosine led to context-dependent induction or inhibition of CD4^+^ T cell differentiation. Specifically, in the presence of exogenous IFN-γ, inosine strongly boosted Th1 differentiation of naïve T cells (Figure 4C) whereas in the absence of IFN-γ, inosine reduced Th1 induction (Figure 4D and Figure S10A). We then dissected the molecular mechanism through which inosine enhanced Th1 differentiation and found that addition of ZM241385, a pharmacological inhibitor of adenosine A_2A_ receptor (A_2A_R) signaling, completely abrogated the effect of inosine (Figure 4C). Moreover, addition of cell permeable cyclic AMP (db-cAMP), a signaling molecule downstream of A_2A_R, restored Th1 differentiation and bypassed the need for inosine. In addition, inhibition of protein kinase A (PKA), a downstream effector molecule of cAMP, similarly negated inosine-driven Th1 differentiation (Figure 4C). Lastly, the inosine-A_2A_R-cAMP-PKA signaling cascade led to phosphorylation of the transcription factor cAMP response element-binding protein (CREB) (Figure 4E), a known transcriptional enhancer of key Th1 differentiation factors such as IL-12 receptor and IFN-γ (Samten et al., 2005; Samten et al., 2008; Yao et al., 2013). Indeed, inosine-dependent upregulation of IL12Rβ2 was also observed (Figure S10B). The effect of inosine was T cell-intrinsic and was not mediated indirectly through DCs because the addition of inosine to naïve T cells that had been activated with anti-CD3/anti-CD28-coated beads also enhanced Th1 differentiation, even in the absence of IFN-γ (Figure S10C). Furthermore, induction of Th1 differentiation and phosphorylation of CREB was absent when A2_A_R-deficient T cells were stimulated with inosine (Figure S10D and S10E). In contrast, bypassing the need for A2_A_R signaling by using db-cAMP increased Th1 differentiation and phosphorylation of CREB in A2_A_R-deficient T cells, confirming that the Th1 promoting effect of inosine is depended on A2_A_R signaling (Figure S10D and S10E). In addition, since pCREB is known to bind to key Th1 target genes, we also confirmed that inosine stimulation led to a sustained upregulation of *Il12rb2* and *Ifng* gene transcription in CD4 T cells (Figure S10F and S10G). Importantly, inosine dose response experiments revealed that the physiological concentrations of inosine observed in sera of *B.p.* but not *C.sp.* monocolonized mice were sufficient to induce Th1 activation (Figure S10H). Since adenosine also binds to the A2_A_R we also measured adenosine levels and found extremely low levels in intestinal contents and, importantly, no differences in serum levels between *B.p.* and *C.sp.* monocolonized mice (Figure S10I), indicating that adenosine could not be mediating the ICB-promoting effects of *B.p*. Furthermore, adenosine dose-response experiments revealed that the levels of adenosine in the serum were insufficient to promote Th1 activation and effector function (Figure S10J). We next wondered if inosine could also directly affect tumor cell survival or susceptibility to T cell-mediated killing. Direct exposure of MC38 tumor cells to inosine *in vitro* did not exert any effects on tumor cell viability (Figure S11A). In addition, pretreatment of MC38 tumor cells prior to co-culture with activated tumor-specific T cells did not promote or inhibit T cell-mediated killing of tumor cells (Figure S11B). These data indicate that the anti-tumor effect of inosine is mediated through T cells.

Combined, these data suggest that the effect of inosine on T cells required sufficient co-stimulation (likely by DCs), IL-12 receptor engagement for Th1 differentiation and IFN-γ production for efficient anti-tumor immunity. Classical dendritic cells (cDCs) were found to be the primary source of IL-12 compared to macrophages (Figure S12A and S12B). Thus, we evaluated the impact of cDCs during cancer and ICB-bacteria co-therapy. To do so, bone marrow (BM) cells from cDC-DTR mice were transferred into lethally γ-irradiated recipient SPF mice to allow for inducible, conditional depletion of cDCs. Following BM reconstitution, mice were treated with antibiotics and then gavaged with a mixture of the three previously identified ICB-promoting bacteria, *B.p.*, *L.j.*, and *O.sp.* Ten weeks after γ-irradiation, mice were subcutaneously injected with MC38 CRC cells and when palpable tumors were established, cDCs were depleted by injection of diptheria toxin followed one day later by anti-CTLA-4 treatment (Figure 4F). Depletion of cDCs led to a significant reduction in intratumoral CD8^+^ and CD4^+^ T cell frequencies and IFN-γ production (Figure 4G-J), which resulted in larger tumors (Figure 4K). Similarly, IFN-γ production and proliferation of splenic CD8^+^ and CD4^+^ T cells were also markedly reduced in cDC-depleted animals (Figure S12C-S12F). Therefore, depletion of cDC strongly reduced the efficacy of bacteria-elicited ICB to reduce established tumors, which indicates the requirement for continuous antigen presentation, IL-12 production and T cell co-stimulation by cDCs for efficient ICB therapy.

### Inosine promotes Th1 immunity and tumoricidal effects in vivo

To confirm whether the inosine-mediated Th1 promoting effect *in vitro* also applied to *in vivo* conditions, GF mice were immunized with ovalbumin in combination with CpG as a co-stimulus. One day later mice received inosine or vehicle only intraperitoneally. Inosine increased the proportions of T-bet^+^, IFN-γ^+^ CD8^+^ and CD4^+^ T cells in the MLN (Figure 5A-C), validating our *in vitro* results. In order to assess whether the effect of inosine also promoted anti-tumor immunity, we challenged GF mice with MC38 tumor cells. Upon palpable tumors, inosine or PBS was given orally or systemically in combination with anti-CTLA-4 treatment and CpG as indicated (Figure 5D). Compared to PBS, inosine led to a reduction of tumor weight and increased anti-tumor immunity irrespective of oral or systemic application routes when given together with anti-CTLA-4 and CpG (Figure 5E-G and Figure S12G and S12H). In contrast, in the absence of CpG as a co-stimulus, inosine increased tumor weight and reduced anti-tumor immunity (Figure 5 E-G and Figure S12G and S12H). These results validate our previous *in vitro* findings demonstrating that the effect of inosine is context dependent based on the amount of co-stimulation present.

**Figure 5:**
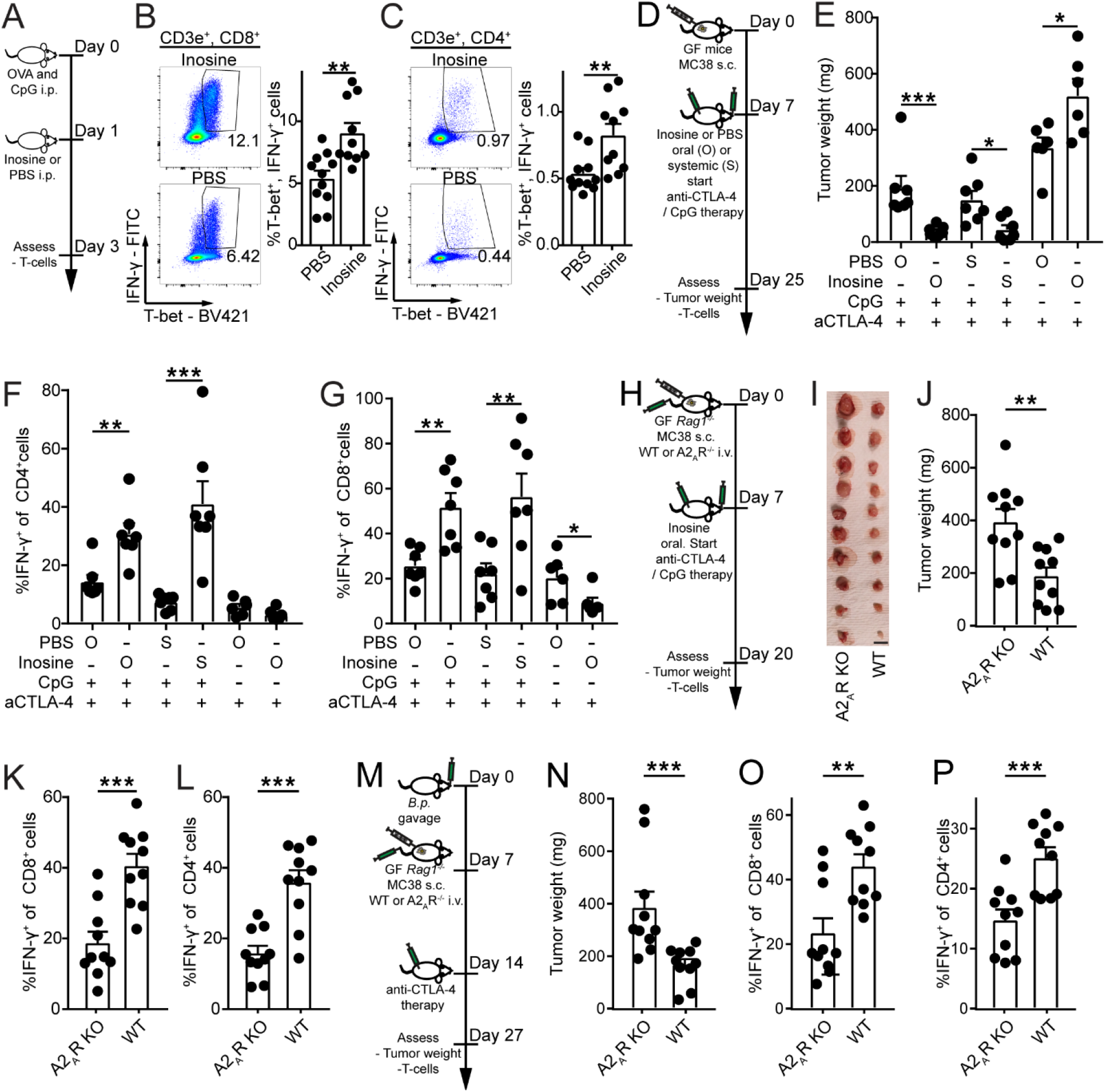
Inosine promotes Th1 activation and anti-tumor immunity. (A) Schematic overview of experimental setup to assess the effect of inosine on Th1 activation *in vivo*. (B and C) Representative dot plots and quantification (left and right panel respectively) of T-bet^+^, IFN-γ^+^, CD3^+^ (B) CD8^+^ or (C) CD4^+^ T cells in the MLN. (D) Schematic overview of experimental setup to assess the effect of inosine on anti-tumor immunity. Upon palpable tumors, mice were treated with 100μg anti-CTLA-4 i.p. (5 times every 72 hours) and in some groups 20μg CpG i.p. (5 times every 72 hours). In addition, inosine (300mg/KG/BW) or PBS was given daily orally (O) through gavage or systemically (S) through i.p. injection. (E) Tumor weight and quantification of IFN-γ^+^ cells amongst (F) CD4^+^ or (G) CD8^+^ cells are shown. (H) Schematic overview to assess the requirement of A2_A_R signaling for inosine-induced anti-tumor immunity. 1×10^6^ MC38 cells (s.c.) and WT or A2_A_R-deficient 1×10^7^ T cells (i.v. 6×10^6^ CD4^+^ and 4×10^6^ CD8^+^ cells) were injected. Upon palpable tumors, mice were treated with 100μg anti-CTLA-4, 20μg CpG (4 times every 72 hours, both i.p.) and inosine (daily, 300mg/KG/BW, through gavage). (I) Pictures of tumors are shown at day 20. scale bars: 1 cm. (J) Tumor weight and quantification of IFN-γ^+^ in (K) CD8^+^ or (L) CD4^+^ cells in the tumor are shown. (M) Schematic overview to assess the requirement of A2_A_R signaling for *B.p.*-induced anti-tumor immunity. Mice were gavaged with B.p. and seven days later 1×10^6^ MC38 cells (s.c.) and WT or A2_A_R-deficient 1×10^7^ T cells (i.v. 6×10^6^ CD4^+^ and 4×10^6^ CD8^+^ cells) were injected. Upon palpable tumors, mice were treated with 100μg anti-CTLA-4 (4 times every 72 hours). (N) Tumor weight and quantification of IFN-γ^+^ in (O) CD8^+^ or (P) CD4^+^ cells in the tumor are shown. Data are mean ± SEM and pooled from two individual experiments (A-P) 10-11 mice/group. *, *P* < 0.05; **, *P* < 0.01; ***, *P* < 0.001. See also Figures S12

We then confirmed that inosine-induced anti-tumor immunity was dependent on A2_A_R signaling *in vivo*. Germ-free *Rag1*-deficient animals were challenged with MC38 tumor cells and simultaneously received WT or A2_A_R-deficient T cells. Seven days later, inosine was given orally in combination with anti-CTLA-4 and CpG (Figure 5H). Inosine enhanced anti-CTLA-4/CpG-mediated anti-tumor immunity in animals that received WT but not A2_A_R-deficient T cells (Figure 5I-L). Specifically, in the presence of inosine, WT but not A2_A_R-deficient T cells displayed increased IFN-γ production within tumors and subsequently only GF mice receiving WT T cells showed reduced tumor weights (Figure 5I-L). Moreover, *B.p.* promoted anti-CTLA-4 efficacy in tumor bearing *Rag1*-deficient mice transfused with WT but not A2_A_R-deficient T cells (Figure 5M-P). This indicated that the ICB-promoting effect of *B.p.* was dependent on A2_A_R signaling and demonstrated a dependency on A2_A_R signaling specifically in T cells for the anti-tumor effect of *B.p.*-inosine-ICB co-therapy.

Taken together, inosine–A2_A_R signaling drives or inhibits anti-tumor immunity *in vivo*, depending on the amount of co-stimulation present.

### Differential effect of ICB-promoting bacteria on CRC subtypes

Lastly, we examined the effect of the identified ICB-promoting bacteria in two distinct models of CRC that mimic different subtypes of human CRC. First, we tested the ICB-promoting effect of *B.p.*, *L.j.*, and *O.sp.* in *Apc^2lox14/+^;Kras^LSL-G12D/+^;Fabpl-Cre* (Haigis et al., 2008) SPF mice, which have conditional *Apc* deficiency and activation of *Kras* specifically in colonocytes. In this model of CRC, anti-CTLA-4 treatment alone did not improve survival compared to isotype-treated animals (Figure 6A and B). To test if the addition of ICB-promoting bacteria could switch a non-responsive to a responsive effect in this model, SPF mice were treated with a mixture of broad-spectrum antibiotics for 7 days to overcome colonization resistance(Lee et al., 2013), followed by bacterial transfer and treatment with anti-CTLA-4. Although the ICB-promoting bacteria colonized the intestine, this combined approach did not enhance survival (Figure 6C and D), revealing a limitation of bacterial co-therapy in this model. Next, we examined the effect of ICB-promoting bacteria in SPF *Msh2^LoxP/LoxP^Villin-Cre* (Kucherlapati et al., 2010) animals that have conditional inactivation of *Msh2* in intestinal epithelial cells. In this model, anti-CTLA-4 treatment alone (without the addition of ICB-promoting bacteria) led to reduced tumor weight and cancer stem cells and increased T cell activation and immune cell infiltration in the tumor (Figure 6E-G, Figure S13A, S13C and S13E). Remarkably, co-treatment with ICB-promoting bacteria boosted the effect of anti-CTLA-4, leading to a further marked reduction of tumor weight and cancer stem cell numbers together with drastically enhanced T cell activation and immune cell infiltration in the tumor compared to control bacteria (Figure 6H-J, Figure S13B, S13D and S13F). In support of a critical role for inosine-dependent upregulation of IL12Rβ2 on T cells and cDC IL-12 production and function, anti-IL-12p75 treatment almost completely abrogated the effect of ICB-promoting, anti-CTLA-4 co-therapy in *Msh2^LoxP/LoxP^Villin-Cre* tumors, which corroborates similar findings upon simultaneous depletion of IL-12 and IL-23, using anti-IL-12p40 treatment (Routy et al., 2018; Vetizou et al., 2015). Finally, since oxaliplatin-anti-PD-L1 co-treatment is a more commonly used therapy in the clinics, we confirmed that ICB-promoting bacteria also promoted the efficacy of oxaliplatin-anti-PD-L1 treatment in SPF *Msh2^LoxP/LoxP^Villin-Cre* (Kucherlapati et al., 2010) animals (Figure S14).

**Figure 6:**
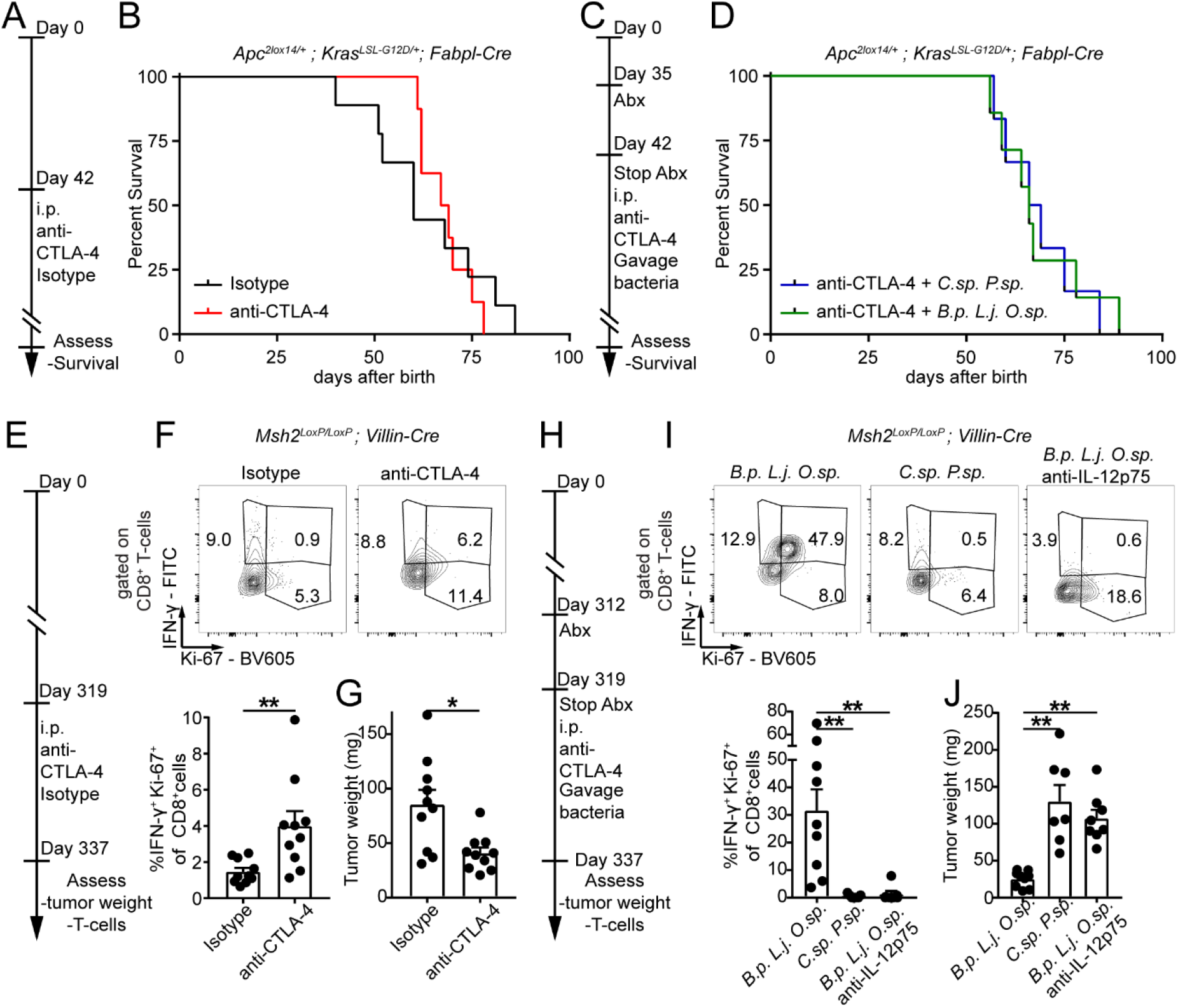
ICB therapy efficacy is CRC subtype dependent. (A, C, E and H) Schematic overview of the experimental setups to assess the effect of ICB-promoting bacteria in different subtypes of CRC. (B and D) Survival curve of isotype, anti-CTLA-4 or ICB-promoting (*B.p. L.j. O.sp*.) or control (*C.sp. P.sp*.) and anti-CTLA-4 co-treated *Apc^2lox14/+^;Kras^LSL-G12D/+^;Fabpl-Cre* animals. (F) Representative plots and quantification of intratumoral IFN-γ ^+^, Ki-67^+^, CD8^+^ T cells and (G) tumor weight of *Msh2^LoxP/LoxP^Villin-Cre* mice. (I and J) same as (F and G), but for bacteria, anti-CTLA-4 co-treated and anti-IL-12p75 co-treated mice as indicated. Data are mean ± SEM and pooled from (B and F) five or (D and I) three individual experiments. (B) *n* = 8-9, (D) *n* = 6-7, (F and G) *n* = 10, (I and J) *n* = 7-9 mice/group. *, *P* < 0.05; **, *P* < 0.01. See also Figures S12, S13 and S14

In summary, our results reveal a novel bacterial-inosine-immune pathway that boosts a cDC-dependent Th1 T cell circuit to greatly enhance the effect of ICB therapies in CRC (Figure S15).

## Discussion

ICB therapy has yielded rather disappointing results in CRC, with an objective response only in 40% of patients with the mismatch repair deficient (MMRD) sub-type of CRC, which amounts to only 4% of all CRC (Le et al., 2015). We have now identified a novel microbial-metabolite-immune circuit that enhances ICB therapy in two mouse models of CRC. These data indicate that modification of the microbiota may provide a promising adjuvant therapy to ICB in CRC. Of note, compared to anti-PD-L1, anti-CTLA-4 induced stronger anti-tumor effects in the AOM/DSS and heterotopic tumor models when both antibodies were administered at the same dose. At this point it is difficult to know if this is due to differences in the biological effects of blocking CTLA-4 versus PD-L1 in these models, but it should be noted that other experimental studies routinely use anti-PD-1 mAb at much higher does than anti-CTLA4 mAb (Routy et al., 2018).

By isolating tumor-associated bacteria we have identified several bacterial species that were found to be associated exclusively with tumors following treatment with ICB, with three of these bacteria able to significantly enhance the efficacy of ICB therapy in CRC. This suggests that the isolation of bacterial species from intestinal tumor biopsies rather than from feces may be a better approach in a clinical setting for defining ICB-promoting bacteria in CRC. Although isolated from mice, all three ICB-promoting bacteria are also found in humans, indicating their potential for translation (Dewhirst et al., 2001; Pridmore et al., 2008; Turroni et al., 2009). Although *Bifidobacterium* species, such as *B. breve* and *B. longum*, have previously been associated with anti-tumor immunity (Sivan et al., 2015), other *Bifidobacterium* species have been reported to provide protection from anti-CTLA-4-induced enterocolitis with no effect on tumor growth(Wang et al., 2018). *Bifidobacterium pseudolongum* species are widely distributed in the mammalian gut with many different strains displaying genomic diversity and differential metabolic capacities (Lugli et al., 2019), suggesting strain-dependent functions and a need for a precision approach to microbial therapy. *Lactobacillus johnsonii* has not previously been associated with anti-tumor immunity; in contrast, it has been shown to have anti-inflammatory effects (Bereswill et al., 2017). Much less is known about the functions of *Olsenella* species.

Our findings demonstrate a critical role for the bacterial metabolite inosine in setting a baseline Th1 level in local mucosal tissues. Initially, this was surprising because previous reports have demonstrated an inhibitory effect of inosine, and A_2A_R engagement in general, on Th1 differentiation *in vitro* and anti-tumor immunity *in vivo* (Csoka et al., 2008; Hasko et al., 2000; He et al., 2017; Ohta et al., 2006). Indeed, the wealth of data supporting an immunosuppressive role for adenosine and A2_A_R signaling has led to the development of novel immune checkpoint inhibitor targets, such as mAb targeting CD73, CD39 and CD38, and pharmacological antagonists of A2_A_R, many of which are currently in clinical trials (reviewed in(Vigano et al., 2019)). However, a small body of literature has demonstrated that inosine can be pro-inflammatory and A2_A_R signaling can sustain Th1/anti-tumor immunity in mice (Cekic and Linden, 2014; Lasek et al., 2015; Lioux et al., 2016). Our findings reconcile these contrasting observations by revealing a context-dependent effect of inosine-A2_A_ receptor signaling based on the amount of co-stimulation. Mechanistically, inosine engages the A2_A_ receptor and activates the transcription factor CREB, through cAMP. CREB, together with co-factors and the formation of heterodimers with ATF-2 and/or c-Jun, modulates the transcription of key Th1 genes, including *Il12rb2* and *Ifng* (Samten et al., 2008). It is worth noting that in addition to cAMP signaling, inosine (compared to adenosine) has a distinct A2_A_R-dependent signaling bias, with a 3.3-fold preference for ERK1/2 phosphorylation. In light of our findings, blockade of inosine-A2_A_ receptor signaling in cancer immunotherapy could negate a positive effect provided by beneficial microbes. We suggest that A2_A_ receptor signaling is likely an integral anti-tumor pathway for bacterial-ICB co-therapies. Indeed, Tanoue *et al* recently identified a consortium of eleven bacteria that improve ICB therapies (Tanoue et al., 2019), which are not related to the bacteria identified in this work. Remarkably though, two of the most elevated metabolites in the cecum and serum of mice colonized with the consortium of 11 bacteria were inosine monophosphate and hypoxanthine, a substrate and product of inosine respectively, which are both A2_A_ receptor agonists like inosine (Welihinda et al., 2016). The identification of this context-dependent effect of inosine-A2_A_ receptor signaling is particularly relevant as inosine is currently used as an intervention in clinical trials in various Th1-associated diseases, including multiple sclerosis, amyotrophic lateral sclerosis and Parkinson’s disease (Bettelli et al., 2004; Kustrimovic et al., 2018; Lovett-Racke et al., 2004; Saresella et al., 2013).

We identified cDCs and their production of IL-12 as essential components for efficient induction of anti-tumor T cell immunity elicited by ICB therapy in the presence of beneficial bacteria. The critical involvement of cDC and IL-12 has also been recently reported upon anti-PD-1 treatment (Garris et al., 2018).

Seminal work by Guinney *et. al* revealed four molecular consensus subtypes of CRC(Guinney et al., 2015), namely MMRD, canonical, metabolic and mesenchymal. In line with the positive results of ICB in MMRD patients in the clinical setting(Le et al., 2015), in our animal model of MMRD (*Msh2^LoxP/LoxP^Villin-Cre*) we indeed observed some efficacy of anti-CTLA-4 single therapy. However, co-therapy with ICB-promoting bacteria strongly enhanced the tumoricidal effect of anti-CTLA-4. Thus, bacterial co-therapy may optimize treatment regimens in MMRD CRC patients. Secondly, ICB therapy was efficacious and was associated with *B. pseudolongum*, *L. johnsonii*, and *Olsenella sp.* in the AOM/DSS model of CRC. AOM/DSS tumors have been used to model inflammation-associated CRC. AOM/DSS tumors also display characteristics of epithelial to mesenchymal transition (Lin et al., 2015), such as reduced E-Cadherin, increased N-Cadherin, Vimentin and SNAIL expression as well as inflammation and increased TGF-β expression (Becker et al., 2004; Mager et al., 2017), which are hallmarks of the mesenchymal consensus molecular CRC subtype (Guinney et al., 2015). Thus, our results indicate a benefit of bacterial co-therapy also in this subtype. Canonical and metabolic CRC subtypes are both characterized by inactivation of *Apc*, canonical additionally by Wnt pathway and metabolic by KRAS activation (Guinney et al., 2015). These hallmarks are well represented in the *Apc^2lox14/+^; Kras^LSL-G12D/+^; Fabpl-Cre* animal model and intriguingly bacterial co-therapy did not improve anti-CTLA-4 treatment. The divergent effect of ICB-promoting bacteria in the *Msh2^LoxP/LoxP^Villin-Cre-* compared to the *Apc^2lox14/+^; Kras^LSL-G12D/+^; Fabpl-Cre* model is intriguing and at this stage we can only speculate about the underlying reason(s). The mutational load and associated number of neoantigens, which is likely higher in *Msh2^LoxP/LoxP^Villin-Cre* tumors, certainly impacts on the efficacy of ICB therapies (Havel et al., 2019). Moreover, anti-CTLA-4 had no effect on its own in the *Apc^2lox14/+^; Kras^LSL-G12D/+^; Fabpl-Cre* model and bacteria alone did not impact on heterotopic tumor development. We also showed that *B.p*. increased the Th1 cell pool and their anti-tumor effect was unleashed followed by effective ICB therapy. Thus, we reason that the discovery of novel checkpoint blockade targets or other therapies that have an effect of their own in the *Apc^2lox14/+^; Kras^LSL-G12D/+^; Fabpl-Cre* model are required to enable efficacious bacterial co-therapy to treat similar subtypes in CRC patients.

Together, this work paves the way for new approaches to precision medicine in CRC.

## Acknowledgements

We are grateful to Carolyn Thomson, Aline Ignacio Silvestre da Silva, Madeleine Wyss, Marcela Davoli-Ferreira, Jenine Yee and Mia Koegler for their help in tackling large scale experiments, their technical knowledge and critical feedback. We thank Mike Dicay for performing the Ussing chamber experiments. Metabolomics data were acquired by RAG at the Calgary Metabolomics Research Facility (CMRF), which is supported by the Canada Foundation for Innovation (JELF 34986) and the International Microbiome Centre (IMC). The IMC is supported by the Cumming School of Medicine, University of Calgary, Western Economic Diversification (WED) and Alberta Economic Development and Trade (AEDT), Canada. LFM was supported by the Early Postdoc.Mobility Fellowship from the Swiss National Science Foundation. KB is supported by a Canada Graduate Scholarship from the National Sciences and Engineering Research Council of Canada (NSERC). MBG is supported by a Canadian Institutes of Health Research (CIHR) grant (PJT-391060) and a Canadian Foundation for Innovation (CFI) grant. KDM is supported by a CIHR grant (PJT-420305), a CFI grant and the Cumming School of Medicine. I.A.L. is supported by an Alberta Innovates Translational Health Chair. JS is supported by a CIHR grant and a Terry Fox Research Institute grant.

## Authors’ contribution

LFM, RB, NC, KB, HR, SP, RAG, IAL, and MG performed experiments and analyzed data. LFM, MBG and KDM wrote the manuscript, and all authors revised the manuscript and approved its final version. JS provided A2_A_R-deficient mice. LFM and KDM conceived the project. KDM and MBG supervised the project.

