## Supplemental Information for "Immunotherapy efficacy in colorectal cancer is dependent on activation of a microbial-metabolite-immune circuit"

*Figure S1 (Related to Figure1)*

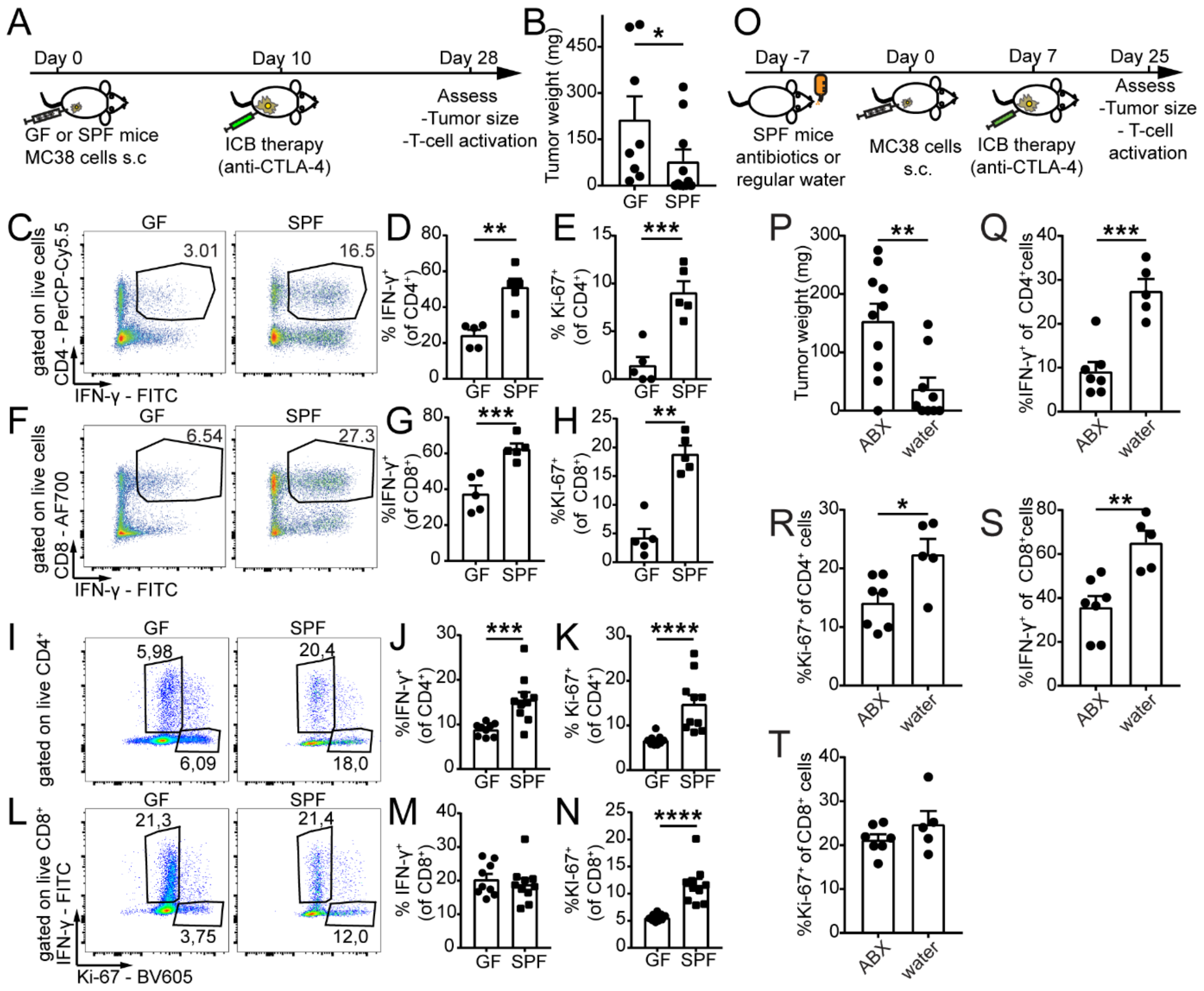


**Figure S1: Bacteria are required for ICB therapy efficacy**. (A) Schematic overview of the experimental setup to determine if bacteria modulate ICB therapy efficacy. (B) Tumor weight at day 28. Intratumoral (C) representative plots and quantification of (D) IFN-$\gamma^+$ and (E) Ki-67$^+$, CD4$^+$ T cells at day 28. (F-G) same as (C-E), but for CD8$^+$ T cells. Splenic (I) representative plots and quantification of (J) IFN-$\gamma^+$ and (K) Ki-67$^+$ CD4$^+$ T cells at day 28. (L-N) same as (I-K), but for CD8$^+$ T cells. Data pooled from two individual experiments ($n$ = 5-10 mice/group). (O) SPF mice were injected with 1x10$^6$ MC38 s.c. and seven days later upon palpable tumors, mice were treated with 100µg anti-CTLA-4 i.p. (5 times every 72 hours). Mice in the antibiotics (ABX) group received a mix of antibiotics (Ampicillin 1mg/ml, Colistin 1mg/ml and Streptomycin 5mg/ml) orally through the drinking water, starting seven days prior to MC38 cell injection until the end of the experiment, whereas mice in the water group received regular water. Tumors were analyzed three days after the last anti-CTLA-4 injection. (P) Tumor weight, quantification of (Q) IFN-$\gamma^+$ and (R) Ki-67$^+$ in CD4$^+$ T cells at day 25 in the tumor tissue. (S and T) same as (Q and R) but CD8$^+$ T cells ($n$ = 9-10 mice/group). Data are mean ± SEM. *, $P < 0.05$; **, $P < 0.01$; ***, $P < 0.001$; ****, $P < 0.0001$.

*Figure S2 (Related to Figure1)*

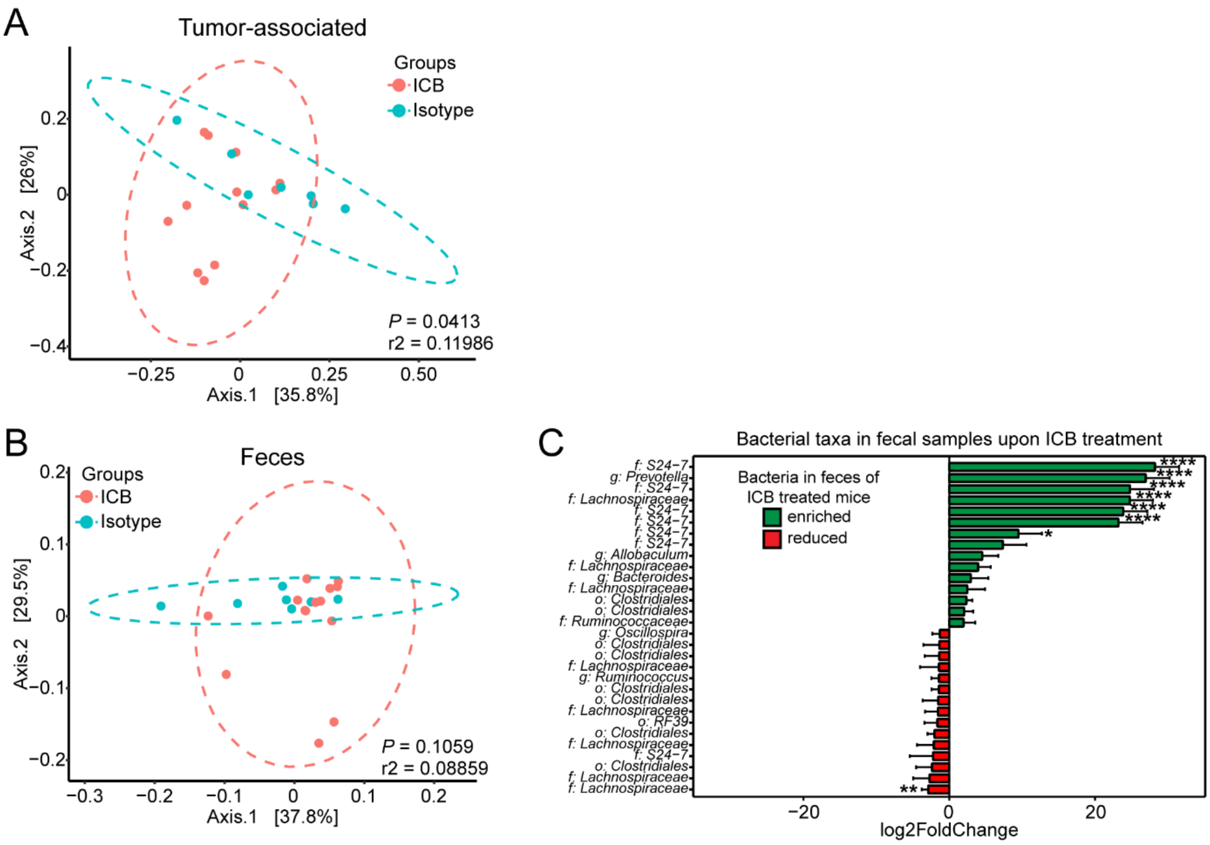


**Figure S2: Microbiota dynamics in ICB treated animals and enrichment in fecal samples following ICB treatment.** (A) Weighted UniFrac PCoA analysis of 16S rRNA gene V4 region amplicon sequencing in tumors of anti-PD-L1/anti-CTLA-4 (ICB) compared to isotype treated animals. (B) same as (A) but for fecal samples. (C) 16S rRNA gene V4 region amplicon sequencing to identify bacteria in fecal samples of mice treated with ICB or control therapies. Bacteria enriched or reduced in fecal samples of anti-PD-L1/anti-CTLA-4 (ICB) compared to isotype treated animals are shown in green or red respectively. Data are mean ± lfcSE (logfoldchangeStandard Error) ($n$ = 7-14 mice/group). Statistics: (A) and (B) PERMANOVA, (C) Benjamini-Hochberg.

*Figure S3 (Related to Figure2)*

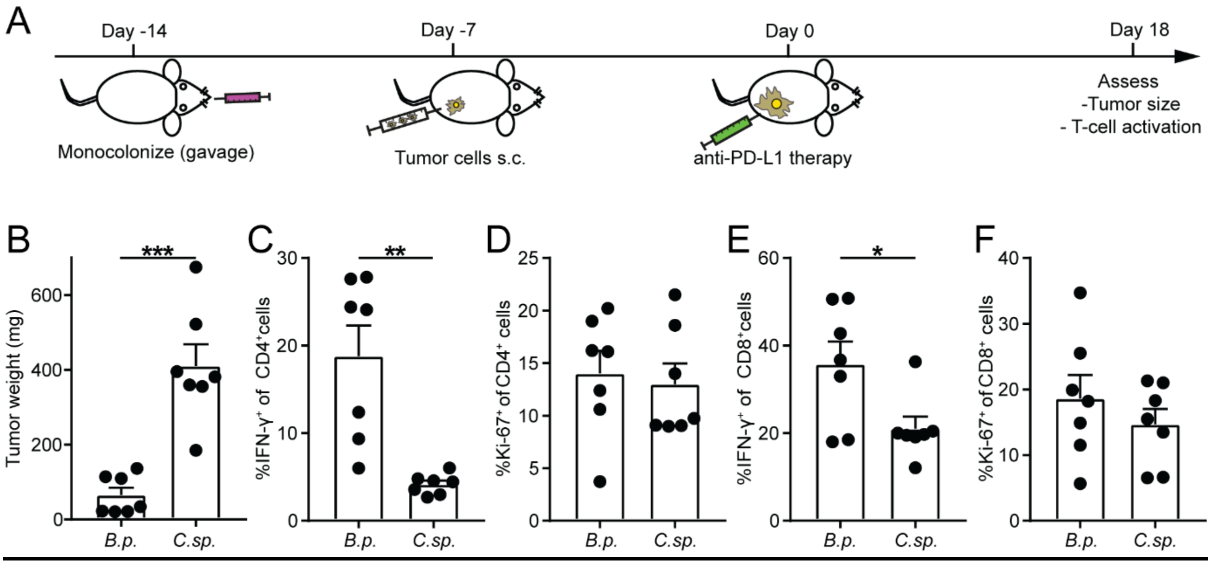


**Figure S3: *B.p.* enhances anti-PD-L1 therapy efficacy**. (A) Germ-free mice were monocolonized with *B.p.* or *C.sp*. Seven days later, 1x10$^{6}$ MC38 cells were injected s.c. and seven days later upon palpable tumors mice were treated with 100µg anti-PD-L1 i.p. (5 times every 72 hours). Tumors were analyzed three days after the last anti-PD-L1 injection. (B) Tumor weight, quantification of (C) IFN-γ$^{+}$ and (D) Ki-67$^{+}$ in CD4$^{+}$ T cells are shown at day 18 in the tumor tissue. (E and F) same as (C and D) but CD8$^{+}$ T cells. Data are mean ± SEM ($n$ = 7 mice/group). *, $P < 0.05$; **, $P < 0.01$; ***, $P < 0.001$.

*Figure S4 (Related to Figure2)*

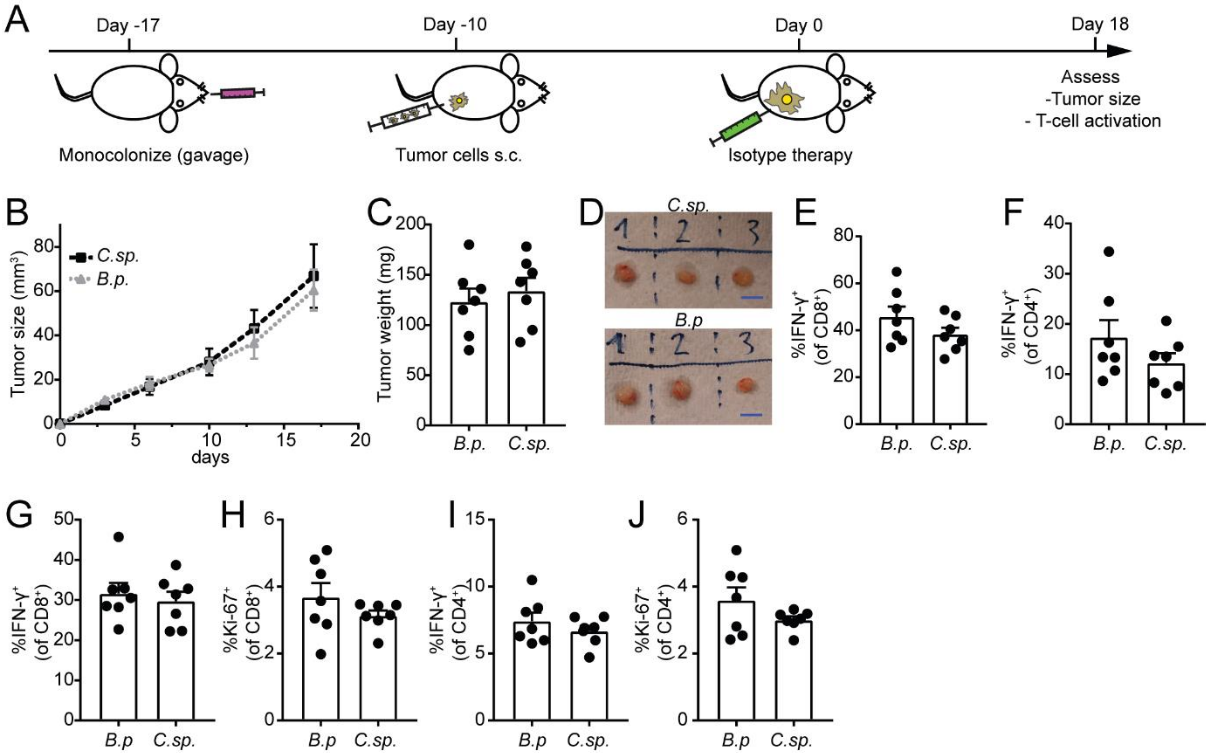


**Figure S4: Bacteria alone to not impact on tumor development.** (A) Overview of the experimental setup to determine whether *B.p.* alone have anti-tumor properties. (B) Tumor growth, (C) tumor weight and (D) representative pictures of tumors are shown, scale bars: 1cm. Intratumoral IFN-γ$^+$ (E) CD8$^+$ or (F) CD4$^+$ T cells at the end of the experiment. Splenic (G) IFN-γ$^+$ or (H) Ki-67$^+$, CD8$^+$ or (I) IFN-γ$^+$ or (J) Ki-67$^+$, CD4$^+$ T cells at the end of the experiment. Data are mean ± SEM ($n$ = 7 mice/group).

*Figure S5 (Related Figure3)*

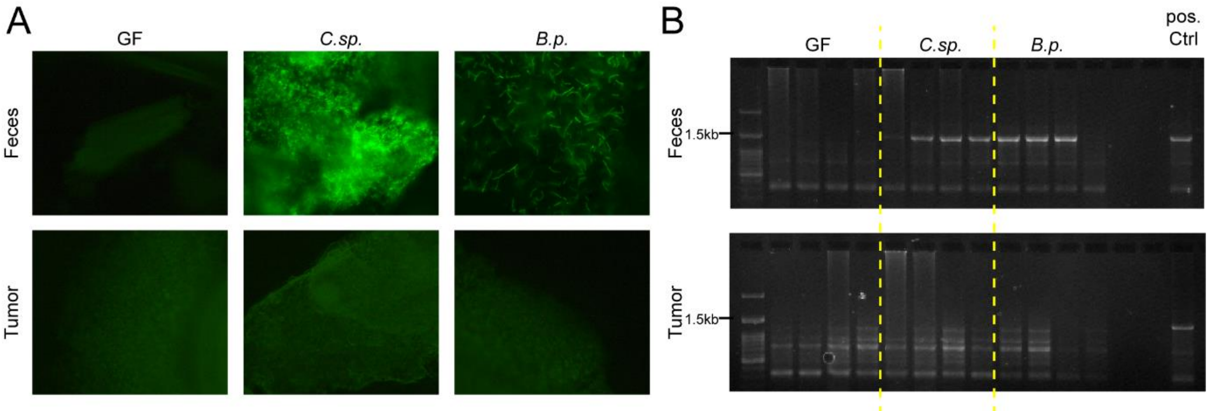


**Figure S5: Bacteria do not translocate into tumor tissue.** (A) Representative pictures of SYTOX green nucleic acid stain of feces and tumor tissue of indicated colonized mice 18 days after initiation of anti-CTLA-4 therapy. (B) Agarose gel of full length 16SrRNA amplicons of feces and tumor tissue of indicated colonized mice.

*Figure S6 (Related to Figure2 and 3)*

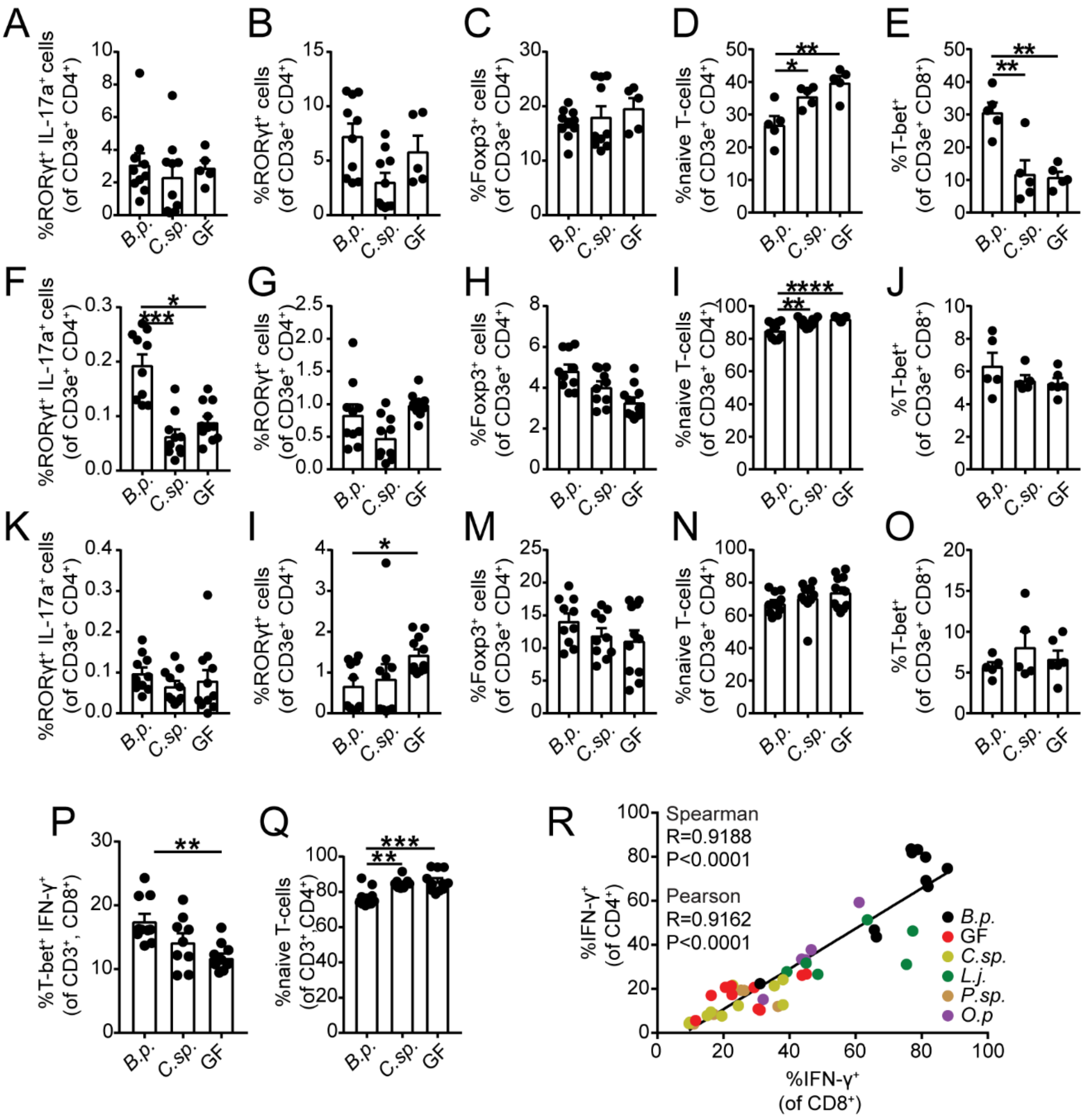


**Figure S6: Effect of *B.p.* on T cell differentiation and activation**. GF animals were monocolonized with either *B.p. C.sp.* or left GF for 28 days before analysis. Small intestinal CD3e$^+$CD4$^+$ T cells expressing (A) RORγt and IL-17a, (B) RORγt, (C) Foxp3 or (D) naïve T cells (defined as RORγt$^-$, GATA3$^-$, Foxp3$^-$, T-bet$^-$). (E) Small intestinal CD3e$^+$CD8$^+$ T cells expressing T-bet. (F-J) same as (A-E), but MLN. (K-O) same as (A-E), but spleen. (P-Q) GF animals were colonized as indicated and after 14 days of colonization treated with anti-CTLA-4 (5 times every 72 hours). Quantification of (P) T-bet$^+$, IFN-γ$^+$, CD3$^+$, CD8$^+$ or (Q) naive (defined as RORγt$^-$GATA3$^-$Foxp3$^-$T-bet$^-$) CD4$^+$ splenic T cells. (R) Correlation of CD8$^+$, IFN-γ$^+$ and CD4$^+$, IFN-γ$^+$ T cells in tumors of anti-CTLA-4 treated, differently colonized mice. Data are mean ± SEM and pooled from two individual experiments (A-Q) $n$ = 5-11 mice/group. (P) $n$ = 46 mice. *, $P < 0.05$; **, $P < 0.01$.

*Figure S7 (Related to Figure3)*

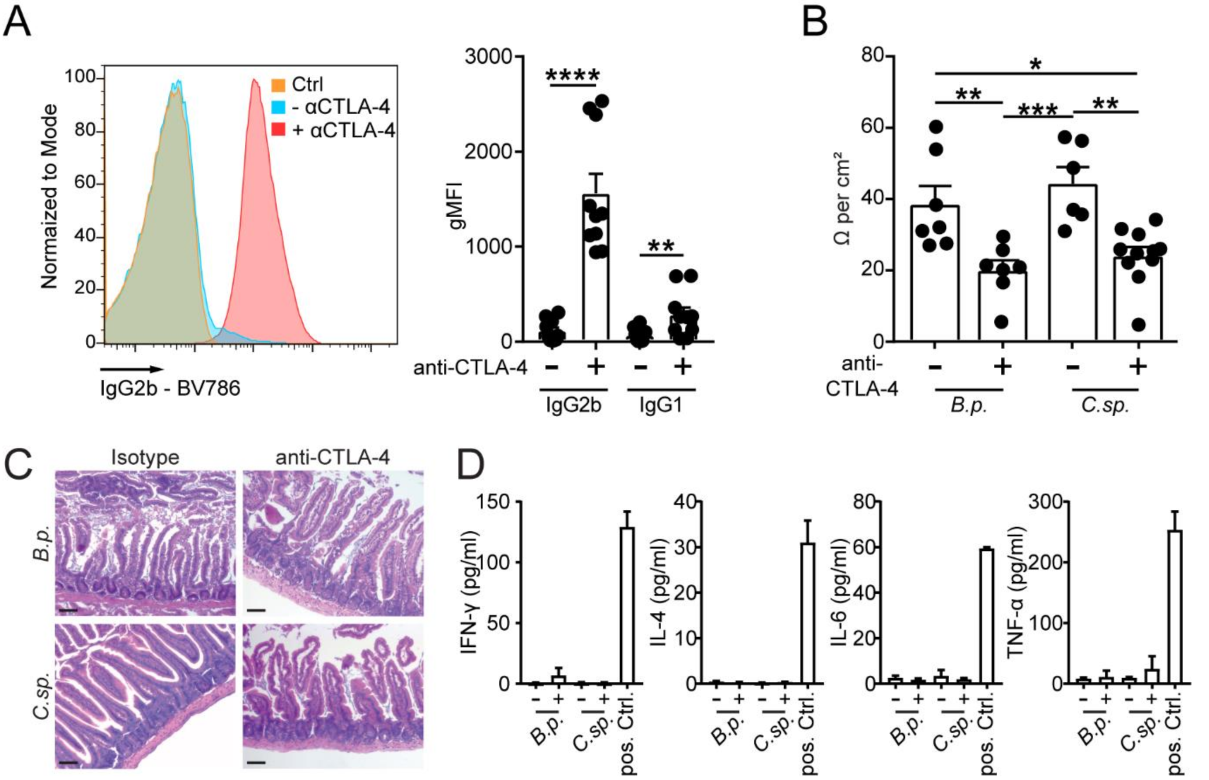


**Figure S7. Reduced barrier integrity upon anti-CTLA-4 treatment**. Serum from monocolonized mice treated with or without anti-CTLA-4 was collected and binding against commensal bacteria was assessed. (A) Systemic IgG2b and IgG1 antibody response upon anti-CTLA-4 treatment in monocolonized mice. (B) Jejunum of B.p. or C.sp. monocolonized mice treated with or without anti-CTLA-4 was collected and barrier integrity was assessed through transepithelial electrical resistance measured in Ussing chambers. (C) Histological inflammation score of small intestinal intestine of *B.p.* or *C.sp.* monocolonized mice treated with or without anti-CTLA-4. Blinded scoring by a board-certified pathologist revealed no inflammation (Scale bar =100µm). (D) Levels of proinflammatory cytokines in the serum of *B.p.* or *C.sp.* monocolonized mice with or without anti-CTLA-4 treatment (100µg i.p. five times every 72 hours) were measured. Serum from DSS-treated SPF mice (2% DSS for 5 days) was used as a positive control for systemic inflammatory cytokines. Data are mean ± SEM and pooled from two individual experiments. (A) $n$ = 9-13 mice/group. (B) $n$ = 6-11 mice/group (C) $n$ = 4 mice/group (D) $n$ = 5 mice/group (pos. ctrl. $n$ = 2 mice). *, $P < 0.05$; **, $P < 0.01$; ***, $P < 0.001$, ****, $P < 0.0001$

*Figure S8 (Related to Figure3)*

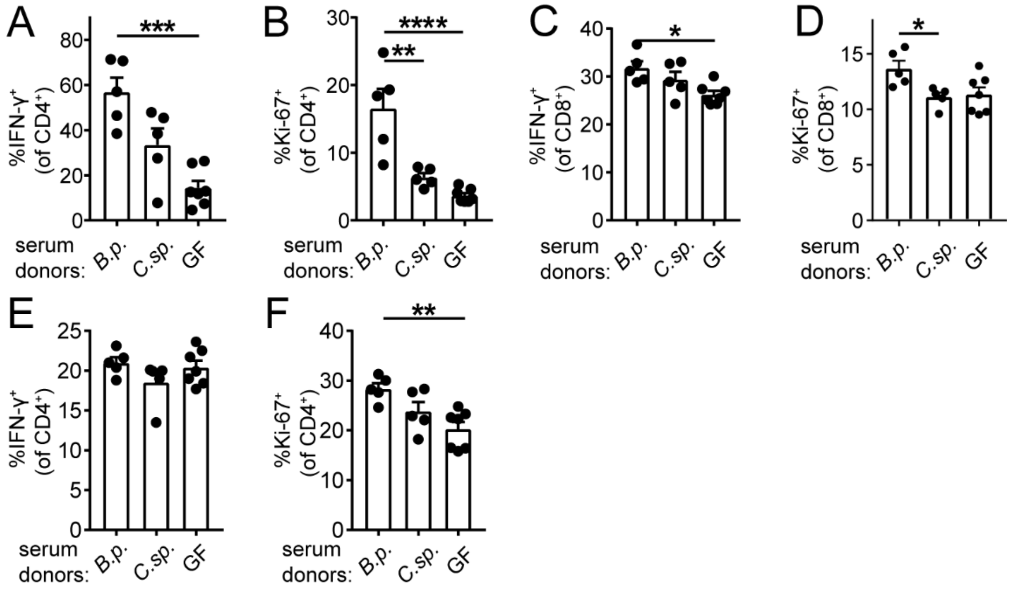


**Figure S8: Systemic anti-tumor immunity upon serum transfer and anti-CTLA-4 treatment.** GF animals were challenged with MC38 tumor cells. Ten days later mice received serum (i.v.) of anti-CTLA-4 treated tumor-bearing animals. Mice were then additionally treated with anti-CTLA-4 (3 times every 72 hours). Serum donors were colonized with *B.p.*, *C.sp.* or remained GF, as indicated. Intratumoral (A) IFN-$\gamma^+$ or (B) Ki-67$^+$, CD4$^+$ T cells. Splenic (C) IFN-$\gamma^+$ or (D) Ki-67$^+$, CD8$^+$ T cells. (E and F) same as (C and D) but CD4$^+$ T cells. Data are mean ± SEM. (A-F) $n$ = 5-8 mice /group. *, $P < 0.05$; **, $P < 0.01$; ***, $P < 0.001$; ****, $P <$ 0.0001.

*Figure S9 (Related to Figure4)*

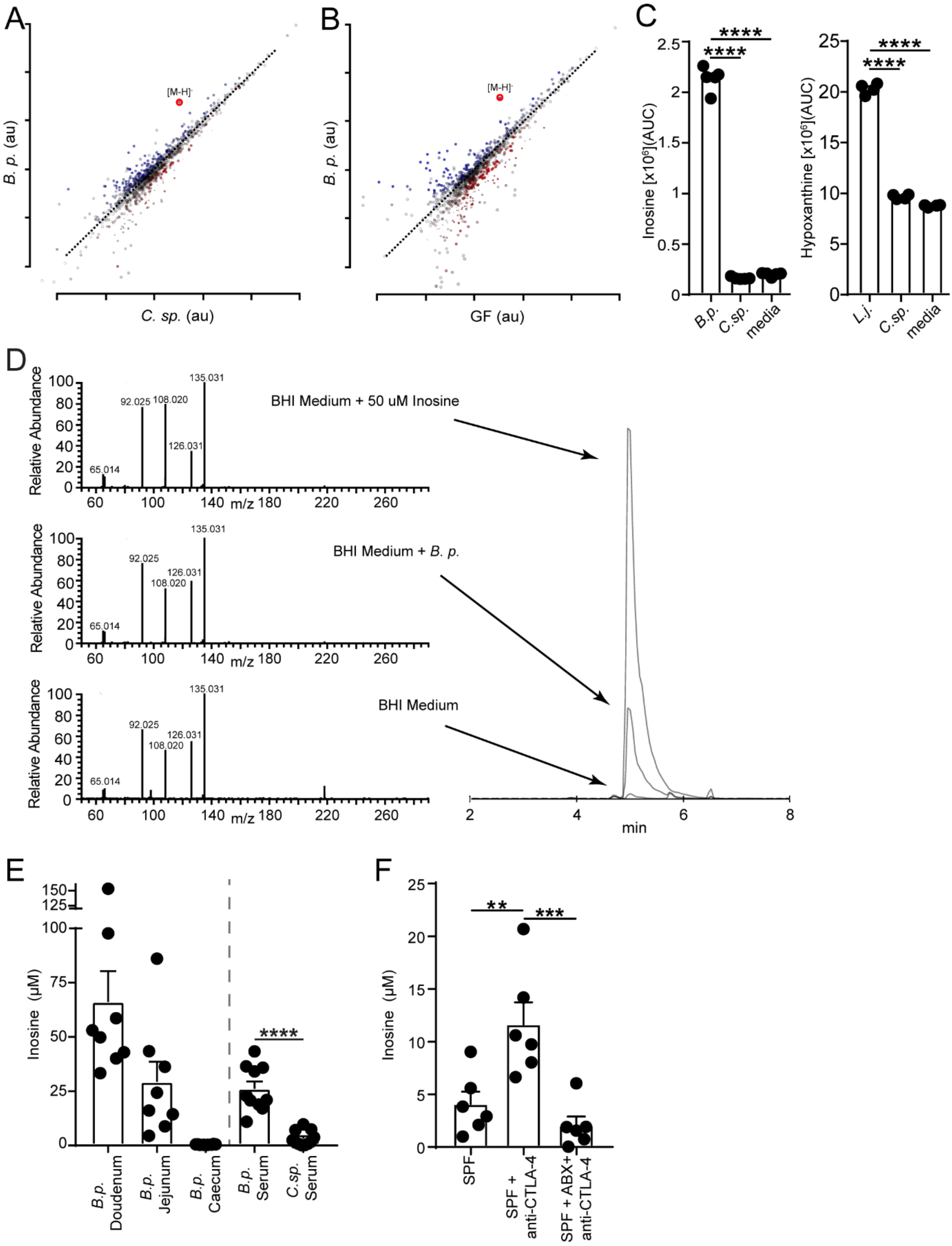


**Figure S9: Inosine levels *in vitro* and *in vivo*.** (A) Scatter plot of untargeted metabolomics data in the serum of anti-CTLA-4 treated, tumor-bearing *B.p.* monocolonized compared to *C.sp*. monocolonized mice. Red circle identifies the inosine signal. (B) Scatter plot of untargeted metabolomics data in the serum of anti-CTLA-4 treated, tumor-bearing *B.p.* monocolonized compared to GF mice. Red circle identifies the inosine signal. (C) Intensity of (left panel) inosine and hypoxanthine (right panel) (AUC: area under the curve) in culture supernatant of indicated bacteria in BHI medium. (D) Parallel reaction monitoring analysis (HCD set at 50eV) of inosine comparing observed fragmentation patterns in BHI medium spiked with and without 50uM inosine as well as *B.p.* cultured in BHI medium. Extracted ion chromatograms of each respective sample are shown in the right panel. (E) Inosine concentrations in duodenal, jejunal or cecal content of *B.p.* monocolonized mice and in the serum of *B.p.* or *C.sp* monocolonized and anti-CTLA-4 treated mice. (F) Inosine concentrations in the serum of untreated (SPF) tumor-bearing, anti-CTLA-4 i.p. (SPF+ anti-CTLA-4) or anti-CTLA-4 plus antibiotic (SPF+ ABX + anti-CTLA-4) treated SPF colonized mice. Anti-CTLA4 treatment: (100µg 5 times every 72 hours). Antibiotics: Ampicillin 1mg/ml, Colistin 1mg/ml and Streptomycin 5mg/ml orally through the drinking water for 32 days. Data are mean ± SEM and pooled from two individual experiments. (C) $n = 5$ biological replicates /group. (E) $n = 8$-11 mice per group. (F) $n = 6$ samples/group. **, $P < 0.01$; ***, $P < 0.001$, ****, $P < 0.0001$.

*Figure S10 (Related to Figure4)*

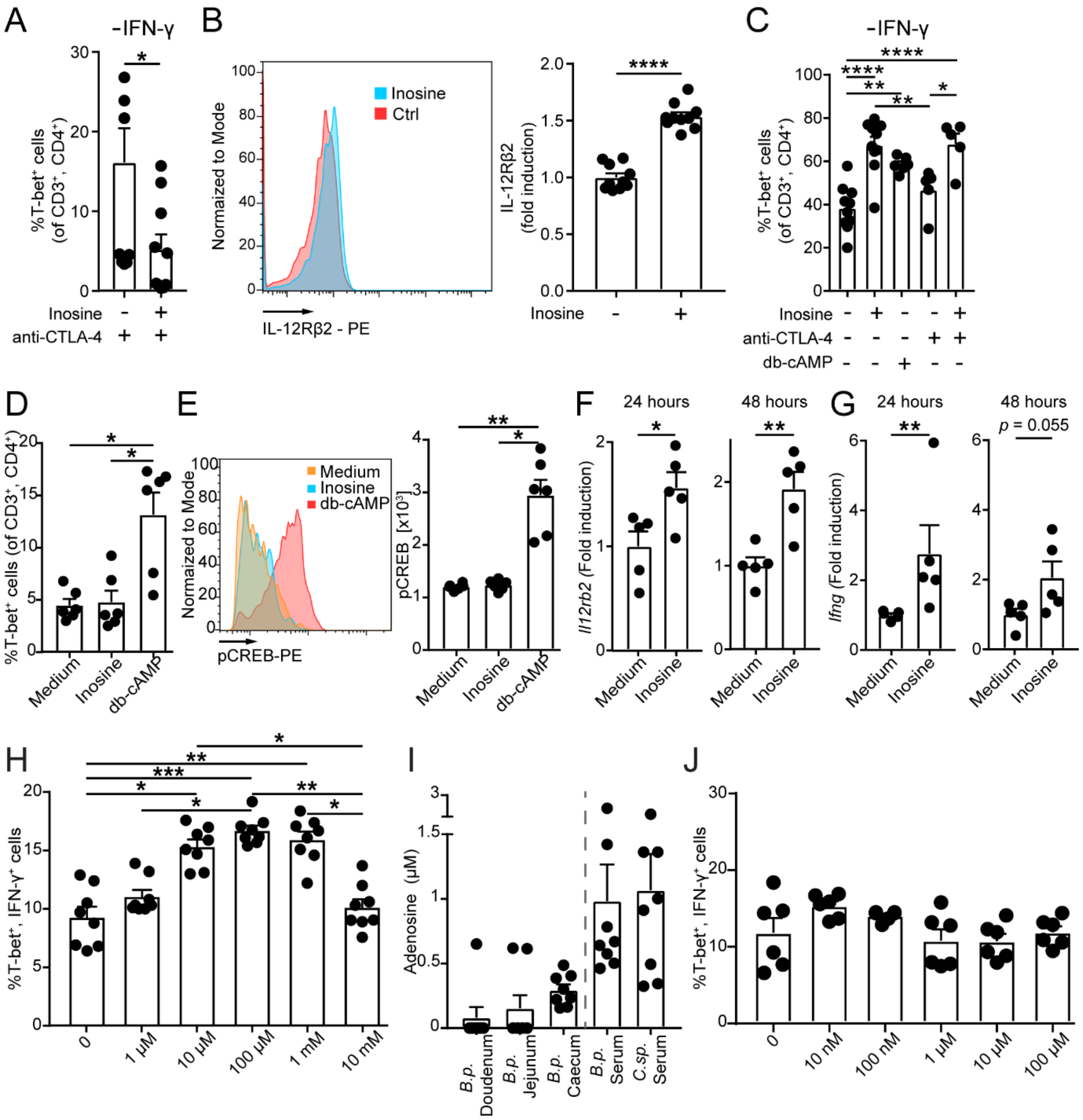


**Figure S10: Context dependent effect of inosine on Th1 T cell differentiation.** (A) Naïve CD4$^{+}$ T cells were co-cultured with bone marrow derived dendritic cells without IFN-γ. Quantification of T-bet$^{+}$, CD3$^{+}$, CD4$^{+}$ T cells 48 hours after co-culture in the presence or absence of inosine and anti-CTLA-4, as indicated. (B) Naïve CD4$^{+}$ T cells were cultured anti-CD3/anti-CD28 coated beads at a ratio of 1:1 without IFN-γ for 48 hours. Representative plot and quantification of IL12Rβ2 surface expression on CD4$^{+}$ T cells in the presence or absence of inosine (left and right panel). (C) Quantification of T-bet$^{+}$, CD3$^{+}$, CD4$^{+}$ T cells 48 hours after co-culture in the presence or absence of inosine, db-cAMP and anti-CTLA-4 as indicated. (D) Naïve, A2AR-deficient CD4$^{+}$ T cells were cultured with anti-CD3/anti-CD28 coated beads at a ratio of 1:1 without IFN-γ. Quantification of T-bet$^{+}$, CD3$^{+}$, CD4$^{+}$ T cells 48 hours after co-culture in the presence or absence of inosine or db-cAMP (E) Representative plot and quantification (left and right panel) of pCREB expression of anti-CD3/anti-CD28 bead co-cultured CD4$^{+}$ T cells in the presence or absence of inosine or db-cAMP 1hour after stimulation. Analysis through flow cytometry. (F and G) Naïve, wild type CD4$^{+}$ T cells were cultured with anti-CD3/anti-CD28 coated beads at a ratio of 1:1. *Il12rb2* and *Ifng* gene transcripts (normalized to *Gapdh*) were evaluated 24- and 48-hours following inosine (1mM) stimulation. Expression was normalized to cells treated with medium. Analysis through quantitative PCR assay (H) Naïve $CD4^+$ T cells were cultured with anti-CD3/anti-CD28 coated beads at a ratio of 1:1 without IFN-γ for 24 hours. Then inosine or vehicle was added at the indicated concentrations for another 48 hours before T cell differentiation and activation was assessed. (I) Adenosine concentrations in the duodenal-, jejunal- or cecal content of *B.p.* monocolonized mice and in the serum of *B.p.* or *C.sp.* monocolonized anti-CTLA-4 treated mice. (J) Naïve $CD4^+$ T cells were cultured with anti-CD3/anti-CD28 coated beads at a ratio of 1:1 without IFN-γ for 24 hours. Adenosine was added then in the indicated concentrations for another 48 hours followed by T cell differentiation and activation as assessed through flow cytometry. Data are mean ± SEM and show pooled data of 2 individual experiments (A and B) $n = 10\text{-}16$, (C) $n = 5\text{-}10$ biological replicates/group, (D and E) $n = 6$ biological replicates/group, (F and G) $n=5$ biological replicates/group, (H) $n = 8$ biological replicates/group, (I) $n = 8$ mice/group, (J) $n = 6$ biological replicates/group. *, $P < 0.05$; **, $P < 0.01$; ****, $P < 0.0001$.

*Figure S11 (Related to Figure4)*

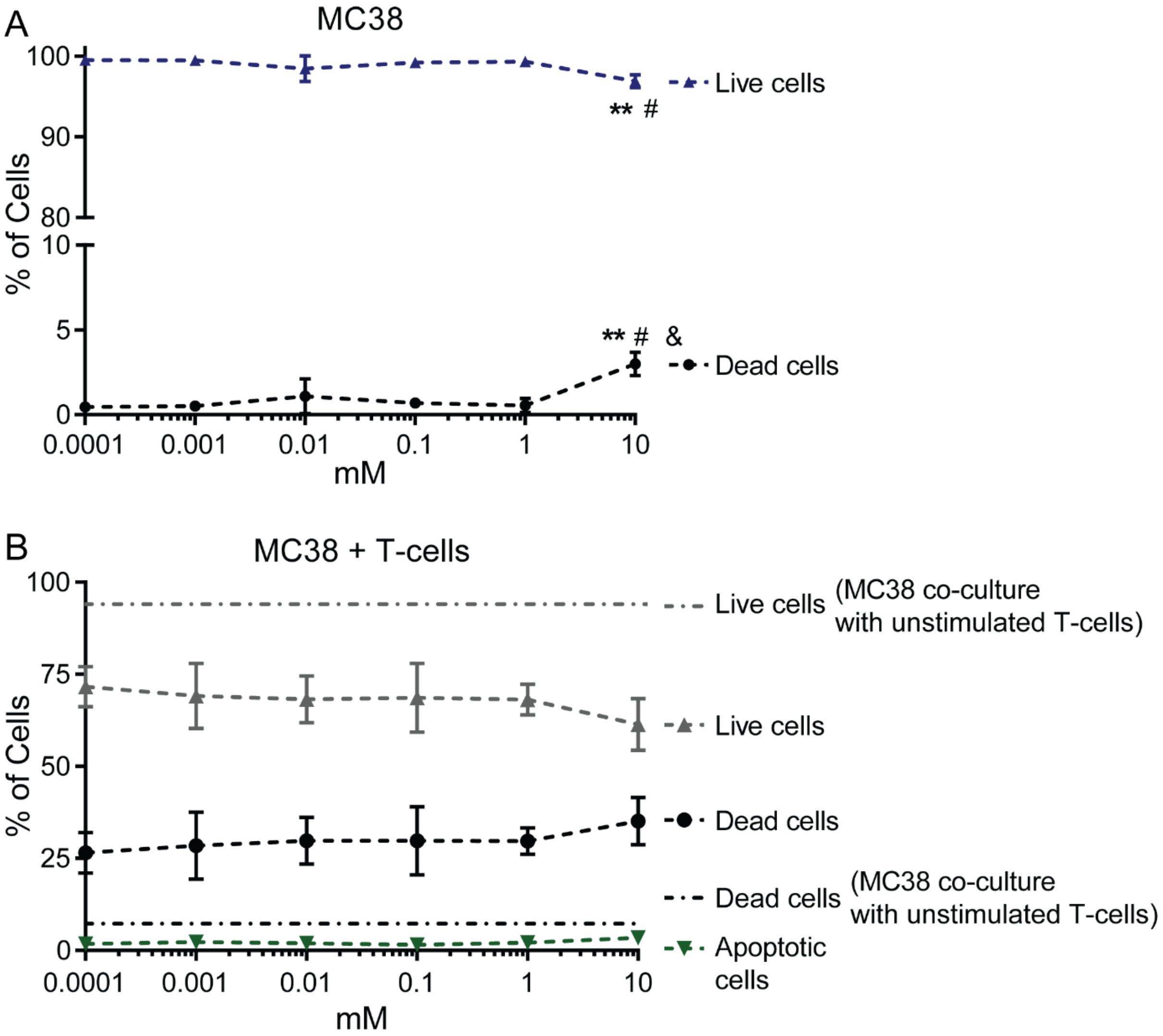


**Figure S11. Inosine does not directly impact tumor cell viability or condition tumor cells for T cell-mediated killing**. (A) MC38 tumor cells were treated with the indicated doses of inosine *in vitro* for 72 hours. Cell death and survival was assessed through flow cytometry. (B) MC38 tumor cells expressing full length ovalbumin (MC38-OVA) were treated with the indicated doses of inosine *in vitro* for 72 hours. In parallel, OVA-specific naïve CD4 and CD8 T cells from spleens of OT-II and OT-I mice, respectively, were activated with anti-CD3/anti-CD28 beads and rmIL-2 (20IU/ml) for 72 hours. Inosine was then washed away from conditioned MC38-OVA cells and fresh medium together with activated T cells were added (100,000 MC38-OVA cells + 25,000 CD4 cells + 25,000 CD8 cells). 72 hours later, cell death and survival of MC38-OVA cells was assessed through flow cytometry. Grey and black dash-dotted lines indicate MC38-OVA cell viability and death when co-cultured with naïve OVA-specific CD4 and CD8 T cells. Data are mean ± SEM and pooled from two individual experiments. $n = 6$ biological replicates/condition. #, & $P < 0.05$, **, $P < 0.01$; (** = 0.0001 vs 10mM, # 0.001 vs 10mM and & 1 vs 10 mM).

*Figure S12 (Related to Figure4 and 5)*

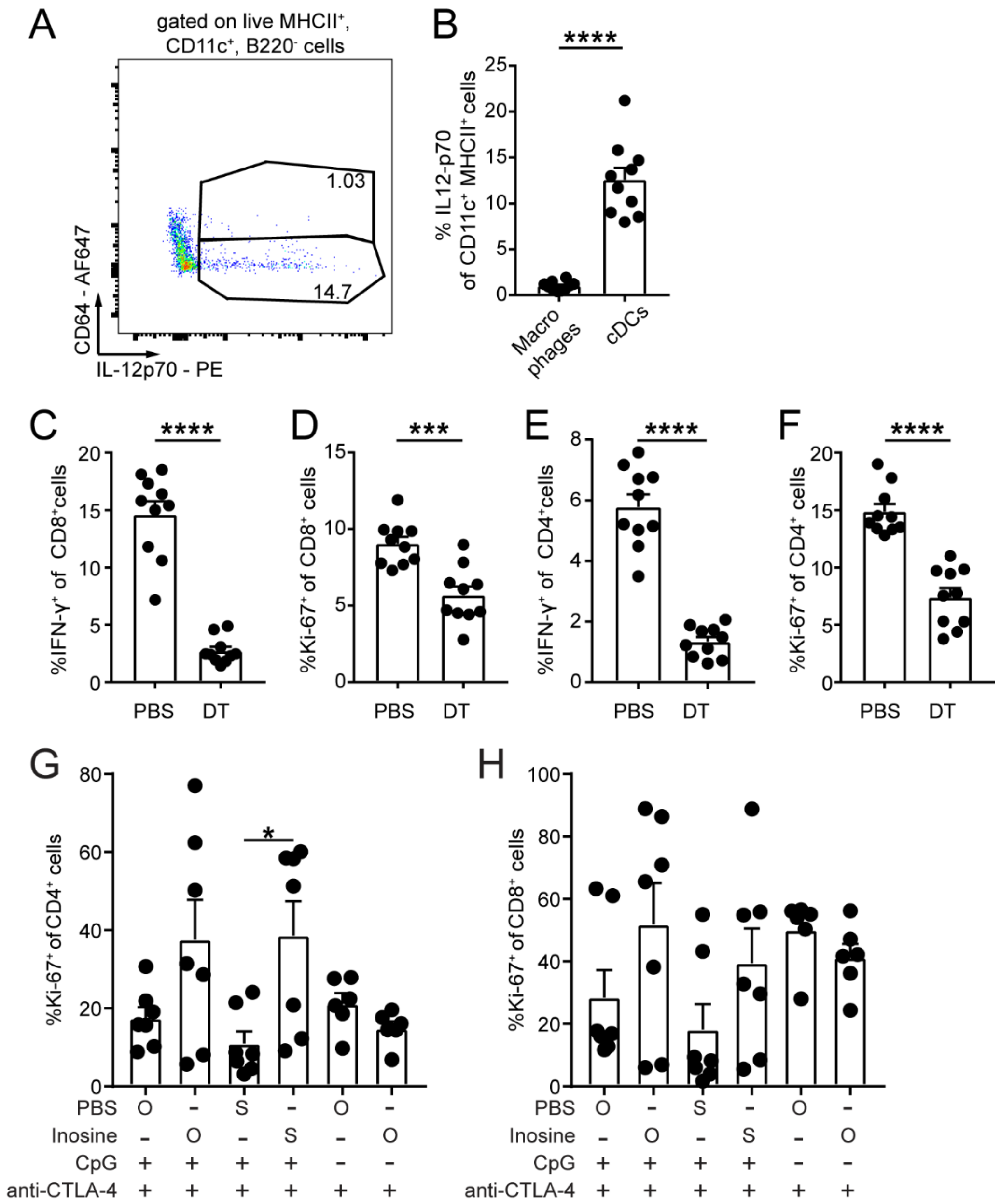


**Figure S12: Classical dendritic cells are required for bacteria dependent effect of ICB.** (A) IL-12p70 expression in classical dendritic cells (MHCII$^{+}$, CD11c$^{+}$, B220$^{-}$, CD64$^{-}$) and macrophages (MHCII$^{+}$, CD11c$^{+}$, B220$^{-}$, CD64$^{+}$) (B) Quantification of IL-12p70 expression in macrophages and cDCs. Classical dendritic cells were depleted with diphtheria toxin (DT) in bone marrow chimeric mice after MC38 tumor challenge, followed by anti-CTLA-4 treatment (see Figure 4H for experimental setup). Quantification of splenic (C) IFN-γ$^{+}$CD8$^{+}$ or (D) Ki-67$^{+}$CD8$^{+}$ T cells. (E and F) same as (C and D) but for CD4$^{+}$ T cells. (G and H) 1x10$^{6}$ MC38 cells (s.c.) were injected in GF mice. Seven days later upon palpable tumors, mice were treated with 100µg anti-CTLA-4 i.p. (5 times every 72 hours) and in some groups 20µg CpG i.p. (5 times every 72 hours). In addition, inosine (300mg/KG/BW) or PBS was given daily orally (O) through gavage or systemically (S) through i.p. injection. Quantification of Ki-67$^{+}$ cells is shown. Data are mean ± SEM and show pooled data of 2 individual experiments. (A - F) $n$ = 10 mice/group. (G - H) $n$ = 6-7 mice/group. *, $P < 0.05$ ***, $P < 0.001$; ****, $P < 0.0001$.

*Figure S13 (Related to Figure6)*

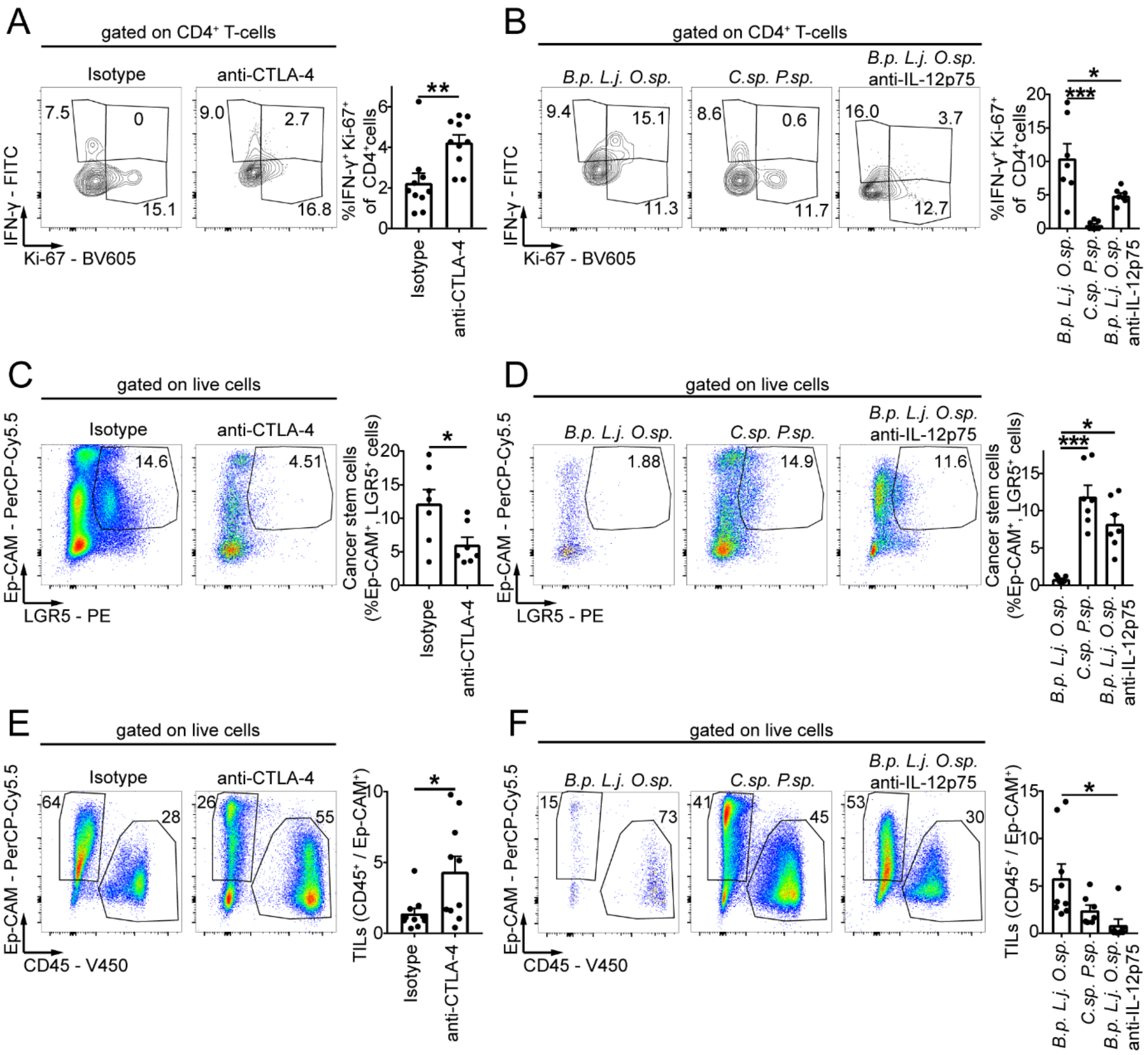


**Figure S13: Bacteria-dependent enhancement of ICB therapy efficacy in *Msh2*$^{LoxP/LoxP}$*Villin-Cre* mice.** *Msh2*$^{LoxP/LoxP}$*Villin-Cre* mice were treated with ant-CTLA-4, ICB-promoting or control-bacteria and/or anti-IL-12p75 (see Figure 5E and H for a detailed experimental setup) (A) Representative plots and quantification of intratumoral IFN-$\gamma^+$, Ki-67$^+$, CD4$^+$ T cells. (B) same as (A), but for bacteria anti-CTLA-4 co-treated and anti-IL-12p75 co-treated animals as indicated. (C) Representative plots and quantification of intratumoral CRC stem cells (defined as Ep-CAM$^+$, LGR5$^+$). (D) same as (C), but for bacteria anti-CTLA-4 co-treated and anti-IL-12p75 co-treated animals as indicated. (E) Representative plots and quantification of tumor infiltrating leukocytes (TILs). (F) same as (E), but for bacteria anti-CTLA-4 co-treated and anti-IL-12p75 co-treated animals as indicated. Data are mean ± SEM and pooled from (A, C and E) five or (B, D and F) three individual experiments. (A, C and E) $n$ = 10 (B, D and F) $n$ = 7-9 mice/group. *, $P < 0.05$; **, $P < 0.01$; ***, $P < 0.001$.

*Figure S14 (Related to Figure6)*

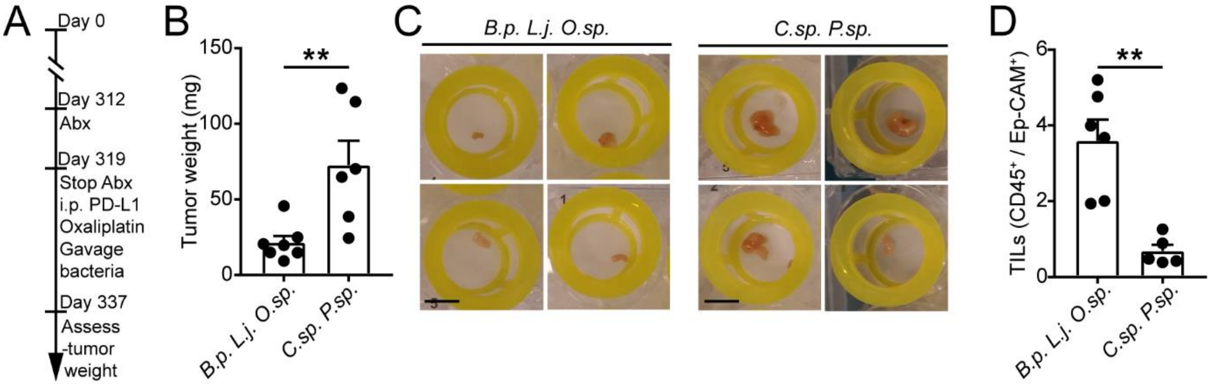


**Figure S14: Oxaliplatin, anti-PD-L1 co-therapy is enhanced by ICB-promoting bacteria.** (A) Schematic overview of the experimental setup to assess the effect of ICB-promoting bacteria in *Msh2*$^{LoxP/LoxP}$ *Villin-Cre* mice. 319 days post birth antibiotics were given orally through the drinking water for seven days (Ampicillin 1mg/ml, Colistin 1mg/ml and Streptomycin 5mg/ml). Then *Msh2*$^{LoxP/LoxP}$ *Villin-Cre* mice were treated with Oxaliplatin, anti-PD-L1 and ICB promoting (*B.p.*, *L.j*. and *O.sp*.) or control bacteria (*C.sp.* and *P.sp*.). Bacteria were given 5 times 72 hours apart through gavage, 100 µg anti-PD-L1 was given 5 times 72 hours apart, i.p. Oxaliplatin 2.5mg/KG/BW was given three times 7 days apart, i.p. (B) Tumor weight of *Msh2*$^{LoxP/LoxP}$*Villin-Cre* mice. (C) Representative pictures of dissected tumors (scale bar: 1 cm). (D) Quantification of tumor-infiltrating leukocytes (TILs). Data are mean ± SEM. *n* = 5-7 mice/group **, *P* < 0.01.

*Figure S15*

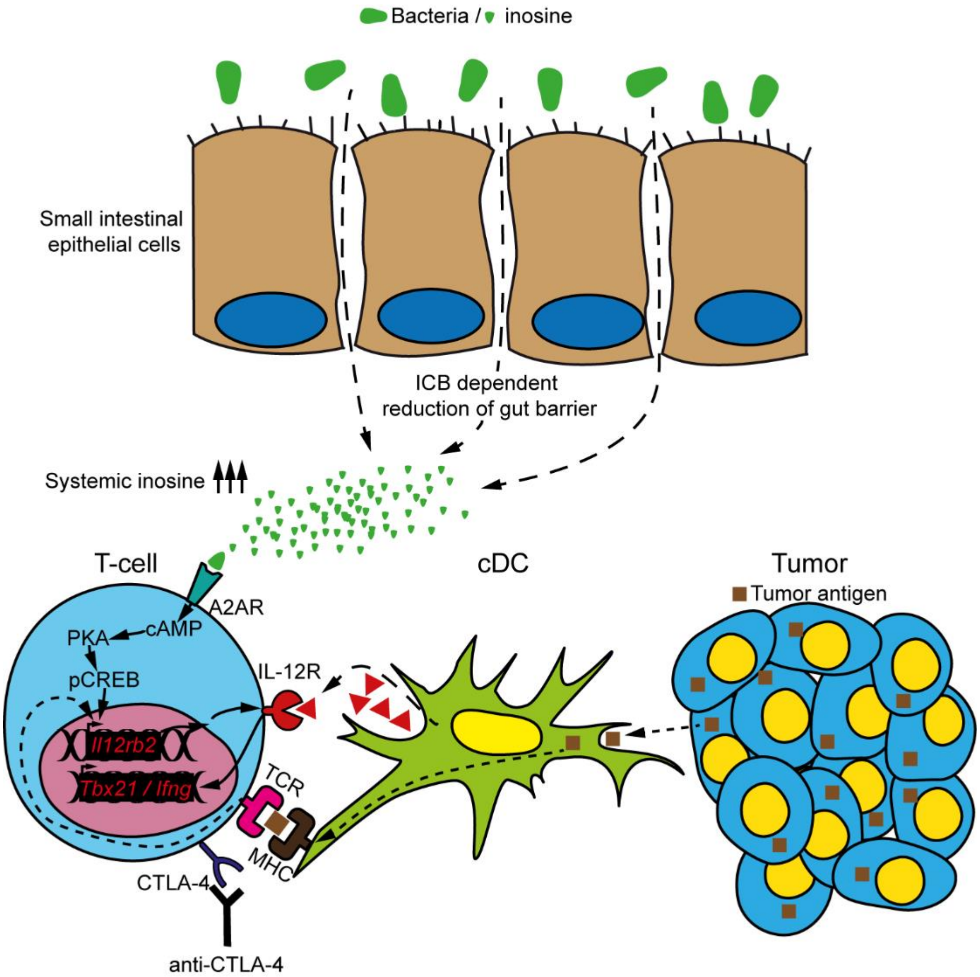


**Figure S15: Mechanism of bacteria-induced ICB efficacy enhancement**. ICB-promoting bacteria increase inosine levels systemically, which is linked to an ICB-dependent reduction in gut barrier integrity. Inosine-mediated $A_{2A}$ receptor engagement leads to increased intracellular cAMP, protein kinase A activation and finally phosphorylation of the transcription factor CREB. Together with TCR stimulation, which is further enabled through anti-CTLA-4 treatment, this leads to increased expression of IL12 receptor on T cells. Classical dendritic cells (cDC) sample antigens and are the major cellular source of IL-12. IL-12 produced by cDCs induces Th1 differentiation, through induction of T-bet (*Tbx21*) expression and activation of T cells. cDCs are required for microbe-anti-CTLA-4 induced IFN-γ (*Ifng*) production by Th1 T cells, which are protective in cancer.

### Material and Methods

*Contact for Reagent and Resource Sharing*

Further information and requests for reagents and resource sharing should be directed to, and will be fulfilled by the Lead Author Kathy D. McCoy.

*EXPERIMENTAL MODEL AND SUBJECT DETAILS*

Animal experiments

C57BL/6J and B6(Cg)-*Zbtb46*$^{tm1(HBEGF)Mnz}$/J (cDC-DTR) (Meredith et al., 2012) and C;129S-*Adora2a*$^{tm1Jfc}$/J mice were obtained from Jackson and then bred and maintained in house. *Apc*$^{2lox14/+}$*;Kras*$^{LSL\text{-}G12D/+}$*;Fabpl-Cre* and *Msh2*$^{LoxP/LoxP}$*Villin-Cre* were kindly provided by Dr. Kevin Haigis and Dr. Winfried Edelman. C57BL/6-Tg(TcraTcrb)1100Mjb/J (Hogquist et al., 1994) (OT-I) mice were bred in house. B6.Cg-Tg(TcraTcrb)425Cbn/J (Barnden et al., 1998) (OT-II) mice were kindly supplied by Dr. Markus Geuking. Global B.6-*Adora2a*$^{tm1Jfc}$/J (Allard et al., 2019) were obtained from Dr. John Stagg (Barnden et al., 1998). All animals were kept in a 12-hour light-dark cycle on standard 4% fat chow. Offspring of different SPF breeding pairs were housed together after weaning to minimize cage effects. Germ free C57BL/6J and *Rag1*$^{-/-}$ mice were bred and maintained in flexible film isolators at the IMC, University of Calgary, Canada. Germ-free status was routinely monitored by culture-dependent and -independent methods and all mice were independently confirmed to be pathogen-free. For experiments, germ-free and monocolonized mice were housed in HEPA filtered Isocages (Tecniplast). Male and female mice between 7-12 weeks were used for experiments, as previous experiments did not show a difference between sexes. In each experiment, mice were age and sex matched and randomly assigned to the different experimental groups. All experiments were performed in accordance with the ethical laws of Alberta and with protocols approved by the Health Sciences Animal Care Committee (AC17-0090 and AC17-0011) following the guidelines set forth by the Canadian Council for Animal Care.

Cell line and primary cell experiments

MC38 parental strain, MC38-EGFP and MC38-OVA female colorectal cancer cells were kindly provided by Dr. Charles Drake. Cells were tested for contamination (Charles River) initially and thereafter screened for absence of mycoplasma every 6-8 weeks (PCR Mycoplasma detection kit, Thermo Scientific). MC38 cell were maintained at 37°C under 5% CO2 in IMDM supplemented with 10% heat-inactivated FBS (Sigma), 100 units/ml penicillin, 100µg/ml streptomycin sulfate, 2mM L-glutamine, 1mM sodium pyruvate and non-essential amino acids (all Thermo Fisher).

Male and female primary bone marrow derived dendritic cells (BMDCs) were generated from the bone marrow of C57BL/6J mice. Cells were maintained in RPMI-1640 (Sigma) supplemented with 10% FBS, 50µm 2-Mercaptoethanol (Sigma) 100 units/ml penicillin, 100 µg/ml streptomycin sulfate and 20ng/ml rm GM-CSF (R&D) at 37°C with 5% CO2.

Male and female naïve $CD4^+$ and $CD8^+$ T-cells were isolated from the spleens of B6.Cg-Tg(TcraTcrb)425Cbn/J, C57BL/6-Tg(TcraTcrb)1100Mjb/J and B.6-*Adora2a*$^{tm1Jfc}$/J mice. Cells were maintained in RPMI-1640 (Sigma) supplemented with 10% FBS, 50µm 2-Mercaptoethanol (Sigma) 100 units/ml penicillin and 100 µg/ml streptomycin sulfate at 37°C with 5% CO2. For BMDCs-T-cell coculture or MC38-Tcell coculture cells were sex matched.

*METHOD DETAILS*

Colorectal cancer models

AOM/DSS tumors were induced in C57BL/6J mice. AOM (10mg/kg/BW) (Sigma) was injected twice at day 0 and 19. 1% DSS (MPBio) solubilized in water was given to the animals 3 times at day 7, 19 and 29 for 5 days followed by regular water for 7 days. (Mager et al., 2017; Mertz et al., 2016). Isotype, anti-CTLA-4 or anti-PD-L1 antibodies (all Bio X Cell) were injected 5 times every 72 hours (100µg/injection) intraperitoneally (i.p.), starting at day 122. *Apc*$^{2lox14/+}$;*Kras*$^{LSL\text{-}G12D/+}$;*Fabpl-Cre* mice have been described previously(Haigis et al., 2008). In short, animals have a median survival of 70 days after birth. Isotype or anti-CTLA-4 antibodies were injected 5 times every 72 hours (100µg /injection) i.p., starting at day 47. In case of microbial transfer co-therapy, antibiotics (ampicillin 1mg/ml (Sigma), Colistin 1mg/ml (Cayman Chemical) and streptomycin 5mg/ml (Sigma), were mixed with water and given *ad libidum* for 7 days starting at day 40 post birth. Bacteria were given through oral and rectal gavage 5 times every 72 hours starting at day 47. Tumor development in *Msh2*$^{LoxP/LoxP}$*Villin-Cre* mice has been described before (Kucherlapati et al., 2010). The median survival of *Msh2*$^{LoxP/LoxP}$*Villin-Cre* animals is 365 days after birth. Therefore, we started treatment with isotype, anti-CTLA-4 or anti-IL12p40 (500 µg, Bio X Cell) antibodies 319 days after birth 5 times every 72 hours. In case of microbial transfer co-therapy antibiotics, same as described above, were given for 7 days starting at day 312 and bacteria were supplied orally through gavage 5 times every 72 hours starting on day 319. For heterotopic cancer models, $1x10^6$ cancer cells were injected subcutaneously (s.c.) in the flank of germ-free, monocolonized or SPF mice. Once tumors were palpable (7-10 days post injection) isotype or anti-CTLA-4 antibodies were injected 5 times every 72 hours (100µg /injection i.p.). For serum transfer experiments, germ-free mice received pooled serum from animals shown in Figure 2. Serum was transferred 3 times (200µl serum each time) every 72 hour intravenously (i.v.). Concomitantly to serum transfer, mice received anti-CTLA-4 three times i.p Tumors were measured every 72 hours using a caliper (length x width x height x π / 6). All tumors were weighed on a fine scale (Mettler Toledo).

Microbiota composition analysis

DNA extraction and purification from feces and cancer epithelial cells was performed using QIAamp Fast DNA Stool extraction kit (Qiagen). The V4 region of the 16S rRNA gene was amplified with barcoded primers (Kozich et al., 2013) using KAPA HiFi polymerase (Roche) under the following cycling conditions: initial denaturation 98°C for 2 min, 25 cycles of 98°C for 30 sec, 55°C for 30 sec, 72°C for 20 sec and final elongation at 72°C for 7 min.

NucleoMag® NGS (Macherey-Nagel) was used for PCR clean-up and size selection followed by PCR product normalization with the SequalPrep™ Normalization Plate Kit (ThermoFisher) according to the manufacturer's protocols. Individual PCR libraries were pooled, then qualitatively and quantitatively assessed on a High Sensitivity D1000 ScreenTape station (Agilent) and on a Qubit fluorometer (ThermoFisher). 16S rRNA v4 gene amplicon sequencing was performed using a V2-500 cycle cartridge (Illumina) on the MiSeq platform (Illumina). Sequences were demultiplexed and processed using the dada2 pipeline (Callahan et al., 2016) within R. Forward and reverse reads were trimmed to 230 and 210 base pairs, respectively. Dereplicated sequences were merged, and chimeras were identified and removed using the removeBimeraDenovo function. Taxonomy was assigned using the Greengenes database (DeSantis et al., 2006). Differentially abundant taxa were identified using DEseq2 (Love et al., 2014), fit to the mean (base mean intensity threshold 20) and with Benjamini-Hochberg correction applied to calculate adjusted p-values. For weighted UniFrac analysis, PERMANOVA was used (9999 permutations).

##### Bacterial culture and identification

AOM/DSS tumors were homogenized in sterile culture media (see below) using a steel bead and a TissueLyser II (Qiagen). Homogenized lysate was streaked on brain heart infusion agar (BHI) and Fastidious Anaerobic Agar (FAA), both supplemented with hemin (5µg/ml), menadione (0.5µg/ml), mucin (250µg/ml), cysteine-HCL (250µg/ml) and Sodium sulfide nonahydrate (250µg/ml) (all reagents from SIGMA) in anaerobe conditions (Whitley, A95 Workstation). Single colonies were picked 48 hours later and cultivated in BHA or FAA medium containing the same supplements as the agar plates. 48 hours after liquid culture, bacteria were lysed and full-length 16S rRNA PCR was performed followed by standard Sanger sequencing. Bacteria were identified by using BLAST. 16S rRNA full length PCR was performed with the following primers: forward1: AGA GTT TGA TCC TGG CTC AG, and forward2: AGA GTT TGA TCA TGG CTC AG together with reverse: ACG GTT ACC TTG TTA CGA CTT (Weisburg et al., 1991).

##### Cell preparation and flow cytometry

Single cells were isolated from spleen, small intestine, mesenteric-, colon draining- and inguinal lymph nodes. Spleen and lymph nodes were cleaned of fat and connective tissue, minced and digested for 20 minutes at 37°C in a shaking incubator (220rpm) in RPMI-1640 supplemented with Collagenase type IA (Sigma) 1mg/ml and DNase I (Roche) 10 IU/ml. Tissues was then filtered through a 40 µm cell strainer (Thermo Fisher) and resuspended in PBS with 2% heat-inactivated fetal bovine serum (FBS) and 2mM EDTA. Fat, connective tissue and Peyer's patches were removed from small intestines, which was then cut into 0.5–1 cm small pieces. Tissue pieces were washed in pre-warmed calcium- and magnesium-free HBSS (Sigma) containing 5 mM EDTA (Sigma) at 37°C in a shaking incubator (220rpm) for 20 min, twice. Supernatant containing intestinal epithelial cells and intraepithelial lymphocytes was discarded. The remaining tissue pieces were then resuspended in in pre-warmed calcium- and magnesium-free HBSS containing Collagenase type VIII (Sigma) 1mg/ml and digested for 20-25 min at 37°C in a shaking incubator (220rpm). Supernatant was filtered first through a 100 µm and then 40 µm cell strainer (Thermo Fisher). For intracellular staining, cells were plated in a 96-well U-bottom plate (Greiner Bio-One) in 200µl IMDM supplemented with 10% FBS, 50 µm 2-Mercaptoethanol, 50ng/ml Phorbol 12-Myristate 13-Acetate (PMA), 750ng/ml Ionomycin and 10µg/ml Brefeldin-A (all Sigma) and incubated at 37°C, 5% $CO_2$ for four hours. Cells were then incubated in Fcγ receptor blocking antibody (BD Biosciences) for 10 min at 4°C followed by surface staining for 25 min at 4°C. For intracellular staining, cells were fixed and permeabilized using the eBioscience™ Foxp3 Fixation/Permeabilization kit (eBioscience) according to the manufacturer's protocol. Cells were then stained with intracellular markers overnight at 4°C. Prior to acquisition, cells were washed and flow cytometry was performed on a FACSCanto (BD Biosciences). Data was analyzed using Flowjo v10.5.3 (Treestar). For a detailed list of antibodies see: Key resource table.

Dendritic cell depletion

For DC depletion experiments, chimeric mouse generation was adapted from previous reports (Mager et al., 2015; Meredith et al., 2012). In short, C57BL/6J mice were lethally irradiated with 1100cGy, split into two sessions of 550cGy each 4 hours apart in a Gamma Cell Exactor 40 (Nordion). Mice were then injected i.v. with $1x10^7$ whole bone marrow from cDC-DTR mice, followed by two weeks of antibiotic treatment in the drinking water (ampicillin 1mg/ml, Colistin 1mg/ml and streptomycin 5mg/ml). Mice then received normal drinking water and were gavaged with a mixture of ICB-promoting bacteria (*B.p., L.j., O.sp.*). $1x10^6$ cancer cells were injected s.c. in the flank 8 week and DC depletion was initiated with diphtheria toxin (100ng every 48 hours i.p., Sigma) 9 weeks after irradiation and bone marrow reconstitution. Anti-CTLA-4 was started one day after the first diphtheria toxin injection and given 5 times every 72 hours. Tumors were measured every 72 hours using a caliper (length x width x height x π / 6) and weighed at the end of the experiment on a fine scale (Mettler Toledo).

Cell culture

Bone marrow derived dendritic cells (BMDCs) were generated from flushed bone marrow cells, maintained in RPMI-1640 (Sigma) supplemented with 10% FBS, 50µm 2-Mercaptoethanol (Sigma) 100 units/ml penicillin, 100 µg/ml streptomycin sulfate and 20ng/ml rm GM-CSF (R&D). Medium was exchanged after 48 hours and 72 hours. 5 days after culture, a magnetic cell sorting step was performed to enrich for $CD11c^+$ cells (Miltenyi Biotec). $CD11c^+$ cells were seeded in 96 flat bottom wells and pulsed with 20ng/ml $OVA_{323-339}$ and 100ng/ml LPS (both Sigma) for 18 hours. In some conditions BMDCs were also cultured with 10ng/ml rmIFN-γ (R&D).

Negative selection magnetic cell sorting (Miltenyi Biotec) was used to enrich naïve OT-II $CD4^+$ T cells. Naïve OT-II $CD4^+$ T cells were then co-cultured with BMDCs at a ratio of 2:1

or stimulated with anti-CD3/anti-CD28 T cell activation beads (Thermofisher) at a ratio of 1:1 for 48 hours prior to restimulation with PMA/Ionomycin in the presence of Brefeldin-A and analysis (see Single cell preparation and flow cytometry for details). In some conditions cells were additionally cultured with various combinations of 2μg/ml anti-CTLA-4, 100μM db-cAMP (Sigma), 5μM ZM 241385 (Sigma), 300μM Rp-8-CPT-CAMPS (Cayman Chemical) or 2mM inosine (Sigma) as described previously (He et al., 2017; Yao et al., 2013).

Quantitative PCR

Naïve $CD4^+$ T-cells were MACS-purified (Miltenyi) according to the manufacturer's protocol. RNA was purified using TRI-reagent (Sigma-Aldrich). RNA was transcribed into cDNA using iScript™ (BioRad). PerfeCTa SYBR Green (Quanta Bio) was used to detect the target genes *Il12rb1*, *Ifng*, and *Gapdh* (QIAGEN). Expression levels of genes were normalized to *Gapdh* mRNA, and medium versus inosine stimulated groups were compared applying the $2^{-\Delta\Delta CT}$ method.

Evaluation of intestinal barrier function

Ussing chamber measurements were performed in one small intestinal section (approximately 3cm long) per mouse was collected from the middle of the small intestine, taking care to exclude Peyer's patches. Electrical resistance was measured in 37°C, oxygenized HBSS after approximately 10 to 15 min of equilibration time(Mager et al., 2017).

Anti-commensal serum antibodies were measured against, *B.p.* or *C.sp.* Bacteria were cultured in anaerobe conditions and then diluted to an O.D.600 of 0.07. Bacteria were then inactivated using sodium-azide. Serum from germ-free, *B.p.* or *C.sp.* monocolonized mice, treated with or without anti-CTLA-4 was heat-inactivated at 56°C for 30 minutes and then incubated with bacterial pellets. Fluorescent secondary antibodies against IgG1 and IgG2b were then used to detect systemic antibodies against pure cultured bacteria (Mager et al., 2017). Serum cytokines were measured by Multiplexing LASER Bead Technology (Eve Technologies).

Metabolomic profile assessment

Metabolites in serum or bacterial cultures were extracted in 50% methanol, centrifuged, and the resulting supernatants were diluted into a linear range for mass spectrometry analysis (1:20 final dilution for microbial cultures and 1:50 total dilution for serum). Ultra-high performance liquid chromatography mass spectrometry (UHPLC-MS) data were then acquired on a Q Exactive™ HF Mass Spectrometer (Thermo Scientific) in negative ion full scan mode (50-750m/z) at 240,000 resolution. Metabolites were separated via UHPLC using a binary solvent mixture of 20mM ammonium formate at pH3.0 in LC-MS grade water (Solvent A) and 0.1% formic acid (%v/v) in LC-MS grade acetonitrile (Solvent B) in conjunction with a Syncronis™ column (Thermo Fisher Scientific). Samples were analyzed using a flow rate of 600uL/min using the following gradient: 0-2 mins, 100 %B; 2-7 mins, 100-80 %B; 7-10 mins, 80-5 %B; 10-12 mins, 5% B; 12-13 mins, 5-100 %B; 13-15 mins, 100 %B. For all runs the sample injection volume was 2uL. Metabolite data were analyzed using the XCMS (Gowda et al., 2014; Tautenhahn et al., 2012) and MAVEN software packages (Clasquin et al., 2012; Melamud et al., 2010). Metabolites were identified by matching observed m/z signals (+/-10ppm) and chromatographic retention times to those observed from commercial metabolite standards (Sigma). Inosine assignments, a key metabolite for this study, were confirmed via MS/MS fragmentation patterns using parallel reaction monitoring. These assignments were further validated by spiking inosine standards into microbial extracts to demonstrate co-retention and matching fragmentation patterns between the observed biomarker and a 50 µM inosine standard.

Effect of inosine *in vivo*

To evaluate the effect of inosine on Th1 activation, mice received 30µg CpG, 100mg EndoFit Ovalbumin (both Invivogen) and 2µg $OVA_{323-339}$ (Sigma) i.p. and 24 hours later mice received 300mg/kg/BW inosine or PBS as a control through i.p. injection. T cell differentiation was assessed 48 hours later. To assess the impact of inosine on tumor development during ICB therapy $1x10^6$ cancer cells were injected subcutaneously (s.c.) in the flank of germ-free mice. Once tumors were palpable (7-10 days post injection) 100µg anti-CTLA-4 antibodies and 20 µg CpG were injected 5 times every 72 hours injection i.p.. 24 hours following the first anti-CTLA-4 / CpG treatment mice received 300mg/kg/BW inosine daily orally (gavage) or systemically (200µl i.p. and 50 µl s.c.) until the end of the experiment. WT or $A2_A$ deficient cells were isolated from spleens using CD4/CD8 (TIL) MicroBeads (Miltenyi). T-cell purity was >95% and $1x10^7$ T-cells were transferred i.v.

Bacterial detection

SYTOX green nucleic acid stain (Thermo Fisher) was performed according to the manufacturer's instructions. Homogenized feces or tumor tissue was fixed in a 4% paraformaldehyde solution (Sigma) for 30 minutes. Subsequently, samples were diluted in PBS at a ratio 1:5 and stained with SYTOX green nucleic acid stain for 60 minutes prior to picture acquisition (Leica DM2500). 16SrRNA full length PCR was performed as described in above ("Bacterial culture and identification").

Experimental design

Sample size of animal experiments was estimated by power analysis and adjusted to b=0.1 (https://clincalc.com/stats/samplesize.aspx) also based on previous experience with the chosen animal models. Mice were randomly assigned to experimental groups. No data were excluded and details on experiment repetition are given in the respective figure legends.

*QUANTIFICATION AND STATISTICAL ANALYSIS*

Statistical analysis

GraphPad Prism v.5.04 for Windows was used. If variance between groups was similar parametric tests were used, such as standard student t test or one-way ANOVA with Bonferroni post-test, in case of significantly different variance between groups non-parametric tests, such as Mann Whitney U test or Kruskal Wallis with Dunn's post-test, were applied. Two-way ANOVA with Bonferroni post-test was used for tumor growth curves. Survival was analyzed using the Mantel-Cox Log-rank test. Other tests are denoted in the corresponding figure or table legend. Only statistically significant differences are indicated in the figures. For all statistical analyses: *, $P < 0.05$; **, $P < 0.01$; ***, $P < 0.001$; ****, $P < 0.0001$. Exact $P$ values and statistical tests used for each panel are reported in the source data.

*DATA AND SOFTWARE AVAILABILITY*

Data availability

Sequencing data from the V4 region of the 16S rRNA gene of tumor-associated and fecal bacteria is deposited at the BioProject database (BioProject ID:PRJNA528297). http://www.ncbi.nlm.nih.gov/bioproject/528297

#### Supplemental Items

*Table S1 title*

**Composition of intratumoral bacteria in checkpoint blockade- compared to isotype treated animals**

Related to: Figure 1J

*Table S2 title*

**Serum metabolites of differently colonized, checkpoint blockade treated and tumor bearing animals**

Related to: Figure 4A
